# Interconversion of unexpected thiol states affects stability, structure and dynamics in engineered antibody for site-specific conjugation

**DOI:** 10.1101/2020.09.28.317339

**Authors:** C. T. Orozco, M. J. Edgeworth, P. W. A. Devine, A. R. Hines, O. Cornwell, X. Wang, J. J. Phillips, P. Ravn, S. E. Jackson, N. J. Bond

## Abstract

Antibody drug conjugates have become one of the most actively developed classes of drugs in recent years. Their great potential comes from combining the strengths of large and small molecule therapeutics: the exquisite specificity of antibodies and the highly potent nature of cytotoxic compounds. More recently, the approach of engineering antibody drug conjugate scaffolds to achieve highly controlled drug to antibody ratios has focused on substituting or inserting cysteines to facilitate site-specific conjugation. Herein, we characterise an antibody scaffold engineered with an inserted cysteine that formed an unexpected disulfide bridge. A combination of mass spectrometry and biophysical techniques have been used to understand how the additional disulfide bridge forms, interconverts and changes the stability and structural dynamics of the antibody. Insight is gained into the local and global destabilisation associated with the engineering and subsequent disulfide bonded variant that will inform future engineering strategies.

---

Antibody drug conjugates (ADCs) have become very promising therapeutics in oncology by combining the high specificity of a tumour-recognising monoclonal antibody (mAb) with the potency of a chemotherapeutic small molecule (payload)^[1]^. Combining two therapeutic molecules into a single agent reduces the systemic toxicity of small molecule chemotherapy whilst facilitating the use of more potent cytotoxic agents which, if administered alone, would have significant dose limitation due to toxicity^[2-6]^. ADCs represent a huge area of research with currently nine FDA approved ADCs on the market, including five within the past year, and more than 60 ADCs are being clinically evaluated in more than 200 active or recently completed clinical trials^[7]^ (Clinicaltrials.gov). In the first generation of ADCs, the payload was conjugated to lysines, which lead to a distribution in number and position of drugs attached, resulting in variable drug-to-antibody ratios (DARs)^[8]^. The DAR is an important contributor to the therapeutic index: the dose range within which efficacy is achieved with an acceptable safety profile, and must be tightly controlled. To do this, the field has iterated towards the conjugation of payloads to canonical cysteines and then to strategies that enabled site-specific conjugation, including non-natural amino acids^[9-11]^, the use of enzymes such as formylglycine generating enzyme^[12]^, transglutaminase^[13,14]^ and sortase A^[15]^, as well as point mutations to add unpaired cysteines for conjugation, either by substitution^[16]^ or by insertion^[17]^. In particular, the addition of a cysteine near the hinge region of an IgG 1 scaffold has been investigated^[16-18]^, and both the substitution at position 239 in the heavy chain (S239C) and the insertion after that position (C239i) have given promising results^[16,17]^: both cysteines are easily conjugated, provide stability of the payload over time, decrease FcγR binding and do not affect the binding to the neonatal Fc receptor (FcRn), which should ensure a similar half-life to the wild type scaffold^[16,17]^.

During large scale manufacture of several C239i antibodies, we observed that this scaffold exhibits unusual properties not typical of a wild type IgG1. Investigating further, non-reduced peptide mapping, established that the thiol groups of the inserted cysteines adopt three distinguishable chemical states: free thiol (2xSH); capped with cysteine (cysteinylated, 2xCys); or forming an additional disulfide bridge between both C239i residues (iDSB) (**Figure S. 1, Scheme 1**). This additional disulfide bridge was confirmed by tandem mass spectrometry (MS/MS) to be an additional interchain disulfide bond, downstream of the two canonical disulfide bonds that covalently bond IgG 1 heavy chains (**Figure S. 4** and Supporting Information (S.I.), sections 1.3 and 4.1). Molecules containing both free thiol and cysteinylated C239i were also observed but typically at a much lower abundance.

The discovery of the additional interchain disulfide bridge was highly unexpected since the *α*-carbons of the amino acids at position 239 on each heavy chain are 17.2 Å apart from each other in the crystal structure (PDB: 3AVE), whilst the distance between the two *α*-carbons of the cysteines involved in a disulfide bridge is 6.4 Å (canonical disulfide bridges in 3AVE). This raises questions regarding how the iDSB forms and the resulting structural changes necessary to accommodate it.

**Scheme 1:**
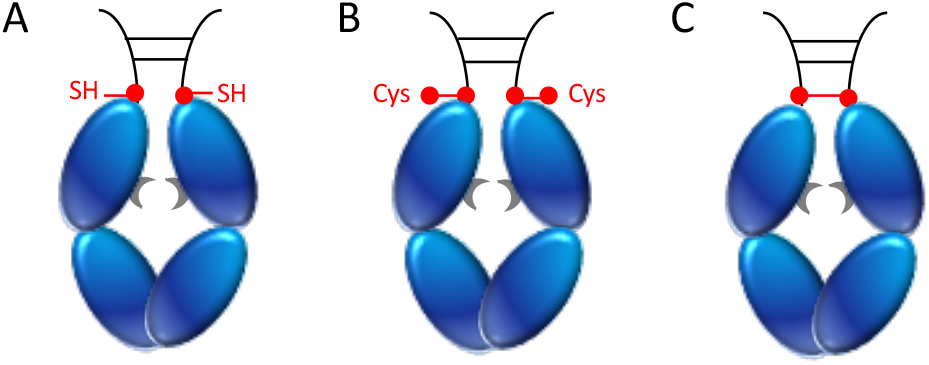
Schematic of the Fc domain containing the inserted cysteine after position 239 (Fc-C239i), in three forms : A. free thiol (2xSH), B. doubly-cysteinylated (2xCys) and C. forming an inter-heavy chain disulfide bond (iDSB). Drawn in grey: glycosylation.

To investigate this systematically, Fc constructs enriched for each of the three thiol states were generated (2xSH, 2xCys and iDSB Fc-C239i), verified, and quantified by mass spectrometry (S.I. sections 1.1.3 and 1.2, **Figure S. 3, Table S. 1)**. The Fc constructs were used as surrogates for C239i IgG since it has been well established that the antibody binding fragment (Fab) does not affect the stability of the Ch2 and C_H_3 domains^[19,20]^. Chemical denaturation experiments, differential scanning calorimetry and millisecond HDX mass spectrometry^[21]^ were used to probe the stability, dynamics and structures of the C_H_2 and C_H_3 domains in each of the enriched states. Together, these studies provide insight into the conditions under which different thiol states interconvert, their relative stabilities and the conformational changes that occur upon formation of an additional disulfide bond.

The evolution of the thiol states was monitored over time using nonreducing peptide mapping (S.I. section 1.4). Under native conditions and incubation for seven days at 25 °C, the relative proportion of all enriched Fc variants did not change significantly **(Figure 1 A, Table S**.2). However, in the presence of chemical denaturant (3.5 M guanidinium chloride (GdmCl), both 2xSH and 2xCys converted into the iDSB Fc-C239i **(Figure 1 A)**. Moreover, slower conversion of 2xCys to iDSB Fc-C239i was observed suggesting that interconversion might not be direct.

**Figure 1:**
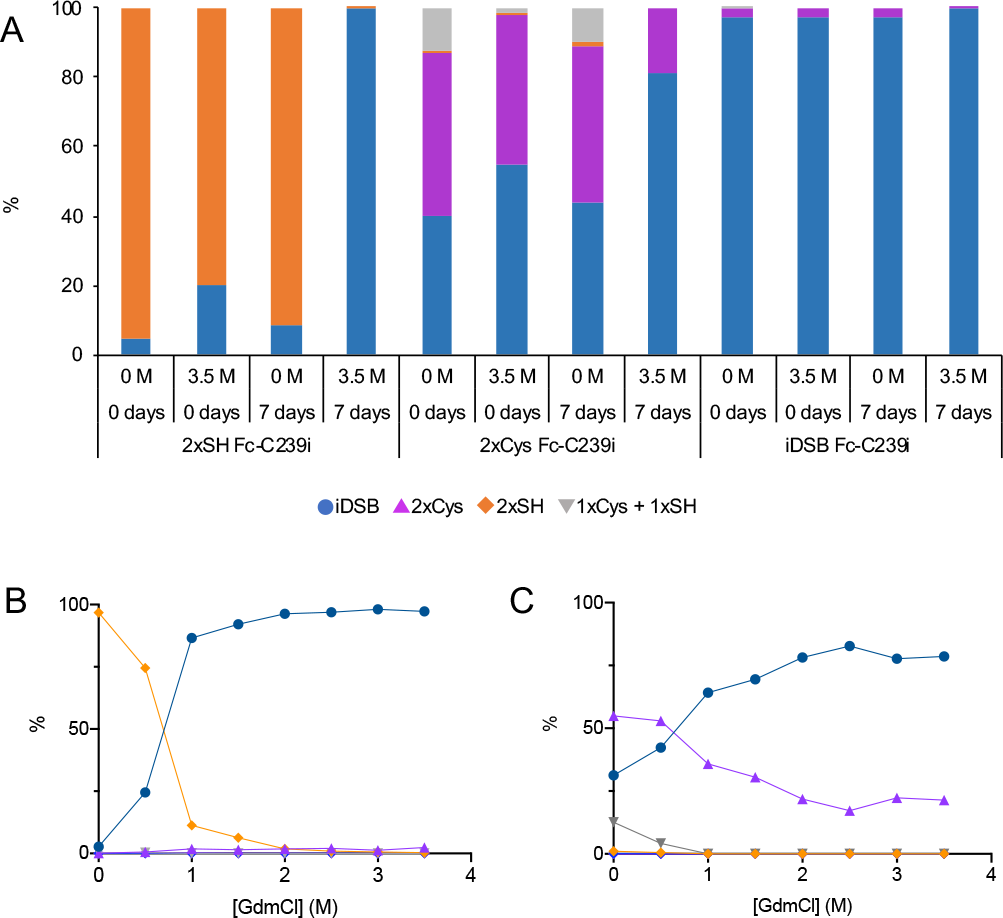
**A**. Effect of the concentration of guanidinium chloride and incubation time at 25 °C in 20 mM His pH 5.5 on the proportion of the thiol states for each of the three initially enriched variants (2xSH, 2xCys and iDSB Fc-C239i). **B**. Effect of the concentration of denaturant after 7 days of incubation at 25 °C on 2xSH Fc-C239i and **C**. 2xCys Fc-C239i enriched starting material. The error bars are too small to be visible.

To understand the interconversion in more detail, the evolution of the thiol states was monitored over a greater range of denaturant concentrations and time points, again by non-reduced peptide mapping (S.I. section 1.5). The conversion from 2xSH to iDSB Fc-C239i was almost complete at denaturant concentrations over 1 M GdmCl **(Figure 1 B, Table S. 4)**. For 2xCys Fc-C239i under equivalent conditions, the conversion to iDSB progressed but not to completion, reinforcing the hypothesis that the doubly cysteinylated form needs to convert into an intermediate, most probably the singly-cysteinylated form, before converting into the iDSB **(Figure 1 C, Table S. 4)**. These data are further supported by the observation that rapid and relatively slow formation of iDSB from 2xSH and 2xCys Fc-C239i, respectively, occur over time at a fixed denaturant concentration (1-4 days at 3.5 M GdmCl, **Figure S. 4)**.

To explain why significant interconversion only occurs in the presence of denaturant (≥ 1 M GdmCl), chemical denaturation unfolding experiments were performed and the thermodynamic stability of both C_H_2 and C_H_3 domains was determined for each of the Fc-C239i enriched variants. The fluorescence derived from solvent exposed tryptophan was measured after seven days of incubation in various concentrations of GdmCl and the data analyzed using the average emission wavelength (AEW, S.I. Equation 1) and fitted to a three-state model (S.I. Equation 3). Three thermodynamic parameters were obtained: 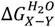 difference in Gibbs free energy between two states X and Y in water; *m*_*X*−*Y*_, the *m*-value between the two states X and Y, a constant of proportionality describing how much the Δ*G*_*X*−*Y*_ changes upon denaturant concentration; and [*den]*_50*%X*−*Y*_, the midpoint of denaturation between states X and Y. All Fc variants showed two unfolding transitions **(Figure 2 A)**, the first corresponding to unfolding of the C_H_2 domain and the second to unfolding of the C_H_3 domain^[22]^. Reproducibility and reversibility were demonstrated (S.I. section 4.2, **Figure S. 6** and **Figure S. 7)**.

**Figure 2:**
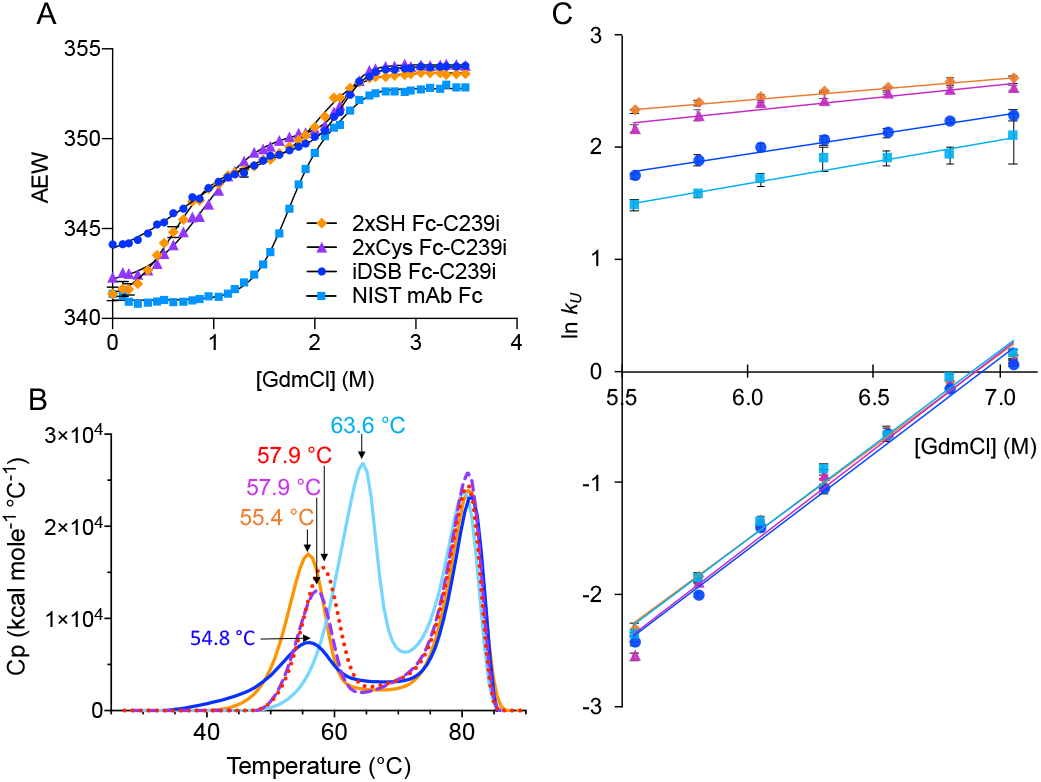
All biophysical experiments on the enriched variants were carried out in 20 mM His pH 5.5. **A**. Unfolding curves of 2xSH Fc-C239i (orange lozenge), 2xCys Fc-C239i (purple triangles), iDSB Fc-C239i (dark blue circles) and NIST mAb Fc (light blue squares). In all cases, the values shown are the average from multiple experiments. **B**. Results from DSC experiments. Thermal stabilities of 2xSH Fc-C239i (orange, solid line), 2xSH NEM-capped Fc-C239i (red, dotted line), 2xCys Fc-C239i (purple, dashed line), iDSB Fc-C239i (dark blue, solid line), NIST mAb Fc (light blue, solid line). The experiment was performed in triplicate but the data shown is only for one experiment. The Tm values shown are the means. For more details on the accuracy on these measurements see S.I. section 4.3. **C**. Unfolding kinetics 2xSH Fc-C239i (orange lozenge), 2xCys Fc-C239i (purple triangles), iDSB Fc-C239i (dark blue circles) and NIST mAb Fc (light blue squares), measured in triplicate. The fastest unfolding phases correspond to the C_H_2 domain unfolding, whereas the slower unfolding phases correspond to the unfolding of the C_H_3 domain. The solid line shows the best fit of the data to Equation 5 (Sup. Info.):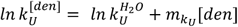. Error bars represent the standard deviation from triplicate measurements. In many cases, they are not visible because they are smaller than the size of the data point.

The midpoint of denaturation observed for C_H_2 domain unfolding in the 2xSH, 2xCys and iDSB Fc-C239i were 0.58, 0.91 and 0.75 M GdmCl, respectively **(Table 1)**. These low denaturation midpoints explain why the conversion from 2xSH and 2xCys to iDSB Fc-C239i starts to occur at 0.5 M and becomes significant over 1 M GdmCl, and suggest the C_H_2 domain has unfolded, at least in part, under these conditions thereby reducing steric constraints and increasing the frequency of disulfide bond formation at the inserted cysteine. A scheme for the interconversion of the different thiol states is shown below, **Scheme 2**, assuming 1xCys + 1xSH can interconvert into iDSB Fc-C239i by nucleophilic attack ^[23,24]^.

**Scheme 2:**
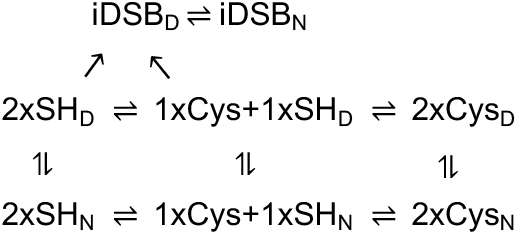
Interconversion network between the 2xSH, 2xCys, iDSB Fc-C239i variants. D: denatured. N: native. 2xCys can interconvert into 1xCys+1xSH which can interconvert into 2xSH from either the native or denatured states, and both 2xSH and 1xCys+1xSH can interconvert into iDSB via the denatured state. The 1xCys+1xSH form could interconvert to iDSB by nucleophilic attack of the free thiol to the capped cysteine.

**Table 1:** Thermodynamic parameters fitted from the unfolding curves The values are average of duplicate (2xCys Fc-C239i, iDSB Fc-C239i) and triplicate (2xSH Fc-C239i, NIST mAb Fc) experiments and the error is the standard deviation of the repeats. For more details on the accuracy on these measurements see S.I. section 4.2.

| Proteins | $m_{I-N}$<br>(kcal mol<sup>-1</sup> M<sup>-1</sup>) | $[den]_{50\% I-N}$<br>(M) | $\Delta G_{I-N}$<br>(kcal mol<sup>-1</sup>) | $m_{D-I}$<br>(kcal mol<sup>-1</sup> M<sup>-1</sup>) | $[den]_{50\% D-I}$<br>(M) | $\Delta G_{D-I}$<br>(kcal mol<sup>-1</sup>) |
| --- | --- | --- | --- | --- | --- | --- |
| 2xSH<br>Fc-C239i | 2.5 ±<br>0.2 | 0.58 ± 0.02 | 1.5 ± 0.2 | 3.1 ± 0.1 | 2.05 ± 0.04 | 6.4 ± 0.3 |
| 2xCys<br>Fc-C239i | 2.099 ±<br>0.003 | 0.9060 ±<br>0.0001 | 1.902 ±<br>0.002 | 5.3 ± 0.6 | 2.30 ± 0.01 | 12 ± 1 |
| iDSB<br>Fc-C239i | 1.3 ±<br>0.1 | 0.75 ± 0.05 | 1.0 ± 0.1 | 4.5 ± 0.2 | 2.272 ±<br>0.002 | 10.3 ± 0.5 |
| NIST<br>mAb Fc | 3.2 ±<br>0.1 | 1.69 ± 0.02 | 5.45 ± 0.2 | 3.9 ± 0.5 | 2.25 ± 0.04 | 9 ± 1 |

To determine if the thermodynamic stability of the antibody had been affected by insertion of the cysteine or the resultant thiol variants, thermodynamic parameters were measured from the chemical denaturation unfolding experiments. All enriched variants showed lower denaturation midpoints for the first transition compared to the wild type **(Table 1)**, suggesting that the C_H_2 domain was destabilized by the insertion of the cysteine in the upper C_H_2 domain. The 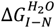 values demonstrate that the C_H_2 domain of iDSB Fc-C239i is the least stable, followed by 2xSH Fc-C239i and finally 2xCys Fc-C239i, which are all significantly less thermodynamically stable than the wild type **(Table 1)**. Since the m-values are correlated with the difference in solvent accessible surface area^[25]^ (Δ*S AA*) between native and denatured states, they provide useful information on the effect of the insertion on the structure. All three variants have a lower *m*_*I*−*N*_-value than the wild type, showing that the C_H_2 domain has a more solvent accessible surface area in the native state suggesting that conformational changes have occurred. Interestingly, the iDSB Fc-C239i has the lowest *m*_*I*−*N*_-value, demonstrating that either the C_H_2 domain has additional solvent exposed surface area in the native state, or that the denatured state is more structured. The midpoint of the second unfolding transition is similar for all constructs, suggesting that the stability of the C_H_3 domain is unaffected by the insertion in the C_H_2 domain.

DSC was employed to measure the thermal stability of the different variants. All constructs showed two unfolding transitions, the first corresponding to unfolding of the C_H_2 domain and the second to the unfolding of the C_H_3 domain **(Figure 2 B)**. The thermal stability of the C_H_2 domain was lower for all Fc-C329i variants than the wild type, and the relative thermal stabilities were the same as those determined using chemical denaturant: iDSB < 2xSH< 2xCys < 2xSH with NEM< NIST mAb Fc **(Figure 2 B, Table S. 7)**. Δ*H*_*cal*_, corresponding to the area under an unfolding peak, is a measure of the favorable interactions that must be overcome to unfold. Δ*H*_*cal*_ of the first transition for the iDSB Fc-C239i was significantly lower than all the other Fc variants and wild type **(Table S. 7)**, indicating that some favorable interactions were lost in the native state of the iDSB Fc-C239i. The peak corresponding to C_H_2 unfolding was also broader for the iDSB Fc-C239i **(Figure 2 B)**, suggestive of a lower Δ*C*_*p*_, which is also correlated to a lower ASASVl^[25]^. This observation is consistent with the *m*-value of iDSB Fc-C239i from the GdmCl experiments suggesting that either the native state of the C_H_2 domain is more solvent accessible than in wild type, or that the denatured state may be more structured. The melting temperatures of the C_H_3 domain for all constructs was very similar, confirming the insertion has no effect on the stability of this domain.

Unfolding kinetics provide information on the rate (and therefore frequency) at which a domain unfolds. Natively folded protein was rapidly diluted into a range of chemical denaturant concentrations and the change in intrinsic fluorescence measured as a proxy for protein unfolding. The data for all Fc variants **(Figure 2 C, Table S. 8)** were fit with a double exponential function (S.I. Equation 4) that best describes both a fast and slower unfolding phase. The unfolding rate constants of the slower phase were very similar for all variants **(Figure 2 C, Table S. 8)** suggesting that this phase corresponds to the unfolding of the C_H_3 domain, and is in agreement with the thermodynamic stability (reported here), as well as the relative kinetic stabilities reported by Sumi and Hamaguchi^[22]^. The unfolding rates of the C_H_2 domain for 2xSH and 2xCys Fc-C239i were very similar, and faster than wild type **(Figure 2 C)**, suggesting that the insertion of a cysteine in the C_H_2 domain kinetically destabilizes it. Interestingly, the rate of unfolding of iDSB was slower than the 2xSH and 2xCys Fc-C239i indicating a higher kinetic stability. This is in contrast to the thermodynamic measurements and is likely due to the unfolding transition state being more structured than for 2xSH and 2xCys Fc-C239i. A more detailed explanation and representation are provided in **Figure S. 9**.

To investigate whether the thermodynamic and kinetic differences associated with the formation of an intramolecular disulfide bond were related to local changes in structure or dynamics, hydrogen-deuterium exchange mass spectrometry (HDX-MS) experiments were conducted. This implementation of HDX-MS uses a novel fully-automated system capable measuring hydrogen-deuterium exchange in the millisecond to minutes time scale^[21]^. Post labelling, online pepsin digestion was conducted and complete sequence coverage achieved in the resulting peptide map (**Figure S. 9**). The insertion of a cysteine at position 239 of the heavy chain (Fc-C239i) changes the structure and/or conformational dynamics of the CH2 domain, which is further, and substantially, disrupted by subsequent formation of an interchain disulfide bond (**Figure 3 A**). Significant increases (t-test, p-value < 0.05) in deuterium exchange consistent with either increased solvent exposure and/or reduced intramolecular hydrogen-bonding were observed in three locations: a small increase in exchange between residues Ser254 and Trp277 that are involved in two β-strands facing each other (strands B and C in Figure 3 B&C); a larger increase in residues between Val279 and Leu306 which are in a β-strand at the edge of the β-sandwich and beginning of the next β-strand (strands D and beginning of E in **Figure 3 B&C**); finally an increase in exchange takes place in the two last β-strands of the CH2 domain (strands F and G, **Figure 3 B&C**), where mixed EX1 and EX2 kinetics are observed for the iDSB (**Figure S. 12**). EX1 exchange kinetics are seldom observed and indicative of a local unfolding event happening. This occurs specifically at strands F and G, suggesting these anti-parallel strands unzip without breaking the surrounding H-bonds of the β-sheet. The H-bonding network involved in the β-strand D has been greatly destabilized and partly broken, and the β-sheet composed of the β-strands A, B and E has also been destabilized but to a lower extent. In previous studies^[26,27]^, it had been observed that the β-strand G in the C_H_2 domain was the most destabilized by other mutations in the C_H_2 domain, demonstrating a lower stability in that specific β-strand. Our data suggest that the additional disulfide bridge at the beginning of the C_H_2 domain applies strain to a larger region, i.e. across the C_H_2 domain, distorting it and opening it up compared to the wild type. The 2xSH and 2xCys Fc-C239i show an increased exchange in the same regions but to a lesser extent. The C_H_3 domain does not show any significant increase in deuterium exchange for any of the enriched variants when compared to the wild type (**Figure 3 A&C**).

**Figure 3:**
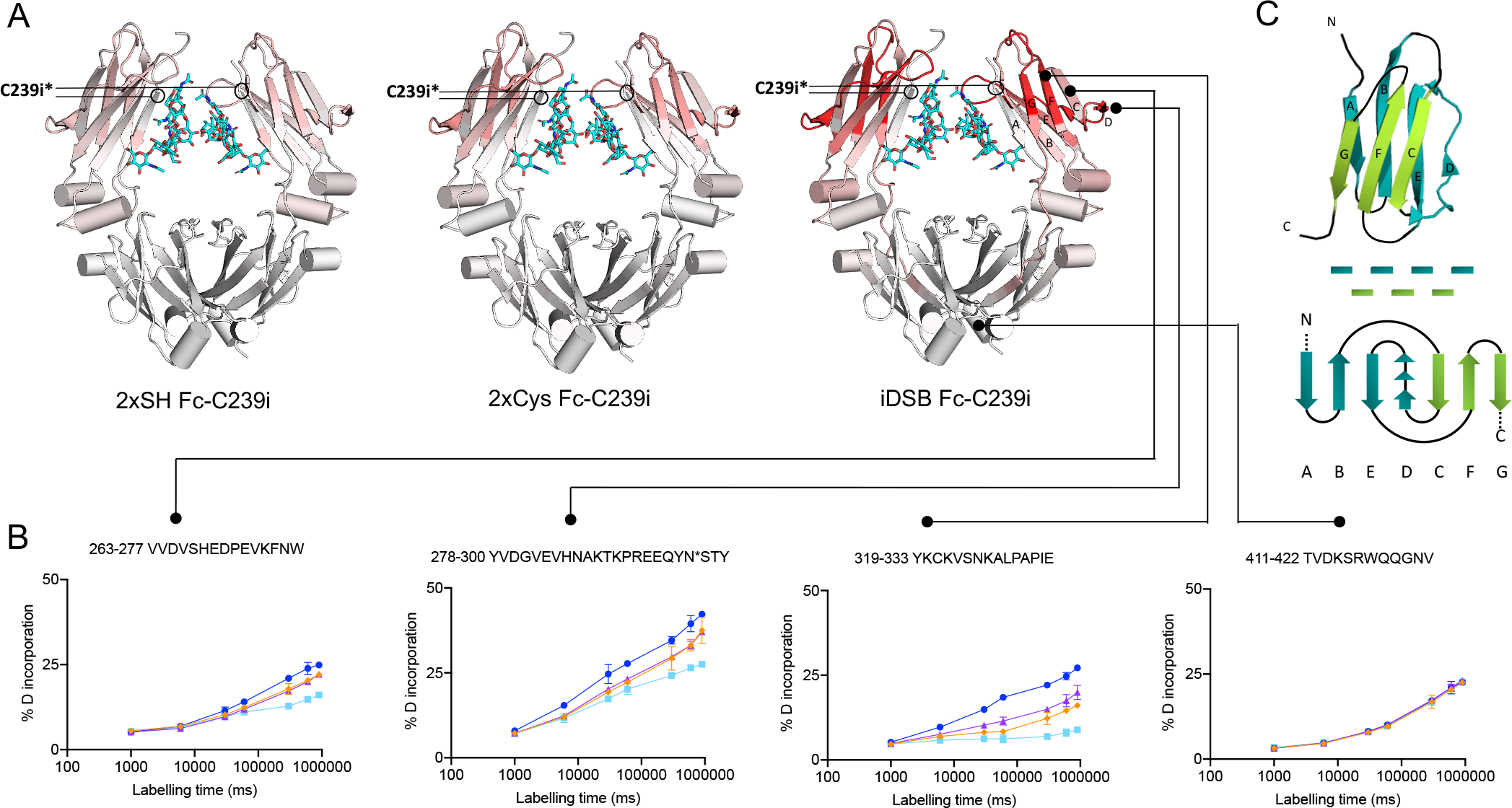
**A**. Crystal structure (3AVE) showing the relative change in fractional deuterium exchange represented as a color scale: reduced exchange (blue), no change (white), increased exchange (red), for the three enriched Fc-C239i variants compared to wild type and normalized to iDSB Fc-C239i. **B**. Uptake plots for different regions of exchange in the Fc domain: 2xSH Fc-C239i (orange lozenge), 2xCys Fc-C239i (purple triangles), iDSB Fc-C239i (dark blue circles) and NIST mAb Fc (light blue squares). The error bars represent a Student’s t-distribution 95% confidence interval, n=3. **C**. Simplified representation of the β-sandwich structure of the C_H_2 domain, indicating the arrangement of the β-strands within the β-sheets and tertiary structure. *The crystal structure 3AVE corresponds to the wild-type and does not have an inserted cysteine at position 239; the annotation shows where the inserted cysteine would be located.

Gallagher *et al*.^[18]^ showed by X-ray crystallography that the inserted cysteine after the 239^th^ residue replaces the position of the serine 239 and affects the preceding residues in the hinge, resulting in a one-residue upward-shift towards the N-terminus of Ser239, Pro238 and Gly237, with residues after the insertion maintaining their wild-type positions. These are rather minimal perturbations that are not consistent with their HDX-MS results. However, our more detailed HDX-MS findings are in agreement with theirs and perhaps reflects that the crystal structure may not represent the true structure in solution due to the restraints imposed by the crystal lattice.

Overall, our studies demonstrate that the C_H_2 domain in all the enriched variants of Fc-C239i has been destabilized by the cysteine insertion, whereas the C_H_3 domain is unaffected. Previous work on antibodies containing either an insertion C239i^[17]^ or the substitution S239C^[16]^ showed that the thermal stability of the C_H_2 domain was not affected by the substitution, but was decreased by the insertion. This shows that the destabilization comes from the insertion, and not from the nature of the residue. In addition to confirming that the insertion does indeed confer a modest but measurable change in structure and dynamics, here we demonstrate a much more substantial structural perturbation can be sustained upon formation of a disulfide bond between inserted cysteines.

Herein we have described, for the first time, an unexpected but naturally occurring state of the C239i mutant containing an additional disulfide bridge, the conditions under which it forms and how it affects the antibody’s structure. More specifically, we showed that the three enriched Fc-C239i variants can interconvert, 2xSH and 2xCys tending towards iDSB. This happens in conditions where the two heavy chains can come close enough to form a disulfide bridge (the sulfur atoms need to be 2 Å apart^[28]^). This can occur in the presence of low concentrations of chemical denaturant but also occurs during expression. When this disulfide bridge is formed, it applies an additional destabilizing strain on the C_H_2 domain. The significant increase in hydrogen deuterium exchange especially in the β-strands C, F, G and in the β-strand D is due to H-bonding breakage and an increase in solvent accessible surface area in the native state. This is also shown by the lower *m*_*I*−*N*_, Δ*Cp* and Δ*H*_*cal*_ values obtained for iDSB compared to 2xSH and 2xCys Fc-C239i. The change in structure of the C_H_2 domain upon forming the additional interchain disulfide bridge might affect not only the native state but also the unfolding transition state, as the unfolding kinetic experiments showed that the iDSB Fc-C239i has greater kinetic stability than the other variants. All these findings suggest that they are caused by a distortion of the C_H_2 domain brought about by the formation of an additional disulfide bridge.

Taken together, these data demonstrate the utility of combining biophysical techniques with high-resolution mass spectrometry to provide detailed characterization of the structure and dynamics of biopharmaceuticals and their variants. Similar observations are anticipated for other engineered antibodies where cysteines are introduced close to the hinge, and the insight provided here serves to guide future cysteine or site-specific engineering strategies.

