## Supplementary Data for "Interconversion of unexpected thiol states affects stability, structure and dynamics in engineered antibody for site-specific conjugation"

### SUPPORTING INFORMATION

### 1 Material and Methods

#### 1.1 Materials

##### 1.1.1 Buffers

Histidine buffer was prepared with the acidic form L-histidine monohydrochloride monohydrate ($C_6H_9N_3O_2.HCl.H_2O$, ≥98% (TLC)) and the basic form L-histidine ($C_6H_9N_3O_2$ ReagentPlus®, ≥99% (TLC)), both from Sigma-Aldrich, USA. The Henderson-Hasselbalch equation was used to calculate how much acid and base were needed to prepare a solution of 20 mM His at pH 5.5. Guanidinium chloride (GdmCl) was purchased from Invitrogen (UltraPure™, ≥99% purity).

For the non-reduced peptide map, the reagents used were: 8 M guanidine hydrochloride solution buffered, pH 8.5 (Sigma Aldrich, USA), N-ethylmaleimide BioUltra, ≥99.0%, HPLC grade (Sigma-Aldrich, USA), EDTA 0.5 M sterile solution biotechnology grade (Amresco, USA), Lysyl Endopeptidase® (Mass Spectrometry grade, Fujifilm Wako Pure Chemical Corporation, Japan), Water for chromatography LC-MS grade LiChrosolv® (Merck, Germany), Acetonitrile for HPLC super gradient Reag. Ph. Eur. USP, ACS water < 30 ppm (VWR, France), Trifluoroacetic acid, Baker HPLC analyzed (Avantor Performance Materials, Poland), sodium phosphate monobasic (meets UPS testing specifications, anhydrous; Sigma-Aldrich, USA), sodium phosphate dibasic (puriss. p.a., ACS reagent, anhydrous, ≥99.0%, Sigma-Aldrich, USA) and sodium chloride NaCl, BioXtra, ≥99.5% (Sigma-Aldrich, USA).

##### 1.1.2 Antibody expression and purification

The antibodies used in this study were NIST mAb Fc, the control protein, and Fc-C239i, a construct described in Dimasi *et al.*[17], with the exact same amino acid sequence as NIST mAb, but with a cysteine inserted at position 239 in the $C_H2$ domain.

All antibodies were produced at AstraZeneca. The gene for the human Fc-C239i (Dimasi *et al.,* 2017) was cloned into a mammalian expression vector. The Fc-C239i protein was transiently expressed in Chinese hamster ovary cells under serum-free conditions. Cleared culture supernatant was loaded directly onto MapSelect SuRe column equilibrated with PBS (pH 7.2). Fc-C239i was eluted with 0.1 M Glycine pH 2.7. Pooled fractions were buffer exchanged into PBS.

##### 1.1.3 Enrichment in the three variants (free-thiol, cysteinylated and interchain disulfide bridge)

The inserted cysteine was shown to be present in three states: free-thiol, doubly-cysteinylated or forming an interchain disulfide bridge. Fc material was enriched in these three states by first diluting the material to 10 mg/mL in 10 mM Tris.HCl, pH 8.0 and reducing the molecule with 2 mM DTT, incubating for 45 minutes at 37 °C. The sample was then buffer exchanged into 10 mM Tris.HCl, using 10 kDa MWCO Amicon spin filters, to remove any remaining DTT and liberated cysteine. For generation of the free thiol (2xSH) variant of Fc-C239i, 1 mM of dehydroascorbic acid (dHAA) was added and incubated for 2 hours at 25 °C. The material was buffer exchanged a final time to remove excess dHAA from the sample. Cysteinylated (2xCys) variants of Fc-C239i were generated through further manipulation of the free thiol variant. L-cysteine (Cys)/L-cystine (Ctn) was added to the sample, at a ratio of 1:4 Cys:Ctn, to a final concentration of 5 mM[29] and incubated at 37 °C for 3 hours. A buffer exchange step was conducted afterwards to remove free L-cysteine/L-cystine from the sample. Disulfide bonded variants (iDSB) were generated by incubating the free thiol variant at 50 °C for 12 hours overnight. The level of enrichment was quantified by targeted mass spectrometry which can chromatographically separate and quantify the iCys states (method in section 1.2, results in Table S.1). The enriched variants were also verified by liquid-

WILEY-VCH

chromatography mass spectrometry and SDS-Page gel (Figure S. 2, Figure S. 3). The liquid chromatography desalting step was carried out on a reverse phase column (Acquity UPLC® BEH300 C4 1.7 µm 2.1 x 50 mm column, Waters Part #186004495), using an aqueous phase A ($H_20$ 0.01% TFA and 0.1% formic acid) and an organic phase B (acetonitrile with 0.01% TFA and 0.1% formic acid) with a two minute gradient from 5 to 80 % B, followed by a three minute gradient from 80 to 95% B. The mass spectrometer (Synapt G2, Waters) was operated in a positive polarity under sensitivity mode, with a capillary voltage of 3.4 kV, source temperature 120 °C, sampling cone at 60 V, desolvation temperature of 400 °C and desolvation gas flow of 800 L/hour.

The free-thiol form was further capped with N-ethyl-maleimide (NEM), by adding 5 µg of N-ethylmaleimide (NEM) to 100 µg of antibody, and incubated at room temperature for 20 min. The excess of NEM was then removed by speed vacuum concentrator.

#### 1.2 Quantitation of the inserted cysteine states

The proportion of the enriched variants was evaluated by targeted non-reduced peptide map.

##### 1.2.1 Non-reduced peptide map preparation: NEM capping, denaturation, and Lys-C digestion

Any free thiols of the antibody samples (25 µg) were capped using 2.5 µg of N-ethyl maleimide (NEM) at room temperature for 20 minutes. Samples were then rotary vacuum evaporated to remove excess NEM. The samples were then reconstituted in 15 µL of a denaturing solution of 7.2 M GdmCl, 5 mM sodium phosphate pH 7.0 and 0.1 M NaCl and was then incubated at 37 °C for 30 minutes. The solution was then diluted by four in 100 mM $NaPO_4$ pH 7.0 containing 0.4 % of 40 mM EDTA, diluting the guanidinium chloride to 1.8 M in preparation for digestion. 0.5 µg of Lys-C were added to 25 µg of thiol capped denatured protein at a 1:50 enzyme:protein ratio and the mixture was incubated at 37 °C for 2 hours. A further 0.5 µg of Lys-C were added and incubated for a final 2 hours. Samples were then analyzed.

##### 1.2.2 Reversed Phase Liquid Chromatography

Mobile Phase A contained 0.1% TFA in Water and Mobile Phase B contained 0.1% TFA in 100% Acetonitrile. The following LC conditions were used: flow rate 0.15 mL/minute, column temperature 55°C throughout the separation with the autosampler maintained at 4 °C. Injections of 10 µL of ~0.42 mg/mL peptide sample were separated using a UPLC Peptide CSH C18, 130Å pore size, 1.7 µm bead size, 2.1 mm X 150 mm column (Acquity). The gradient started at 0% B until a step up to 24% at 4 minutes and then gradually increased to 26% at 8 minutes, followed by a step up to 80%. The gradient was dropped back to 100% A to equilibrated for the next injection. Total run time per samples was 12 minutes.

##### 1.2.3 Mass spectrometry

The Lys-C digested peptides were separated by reversed phase uHPLC coupled to MS (Waters Xevo TQS). The MS desolvation temperature was 600, source cone voltage was 40V, desolvation gas flow 400 L/hr, capillary voltage 3kV, Q3 collision energy 40eV. The different variants were targeted according to the values referenced in the following table.

| Compound | RT (minutes) | *m/z* precursor ion | *m/z* fragment ion |
|---|---|---|---|
| iDSB | 5.70 +/- 0.20 | 1416.50 | 566.30 |
| 2xCys | 5.50 +/- 0.20 | 1476.70 | 566.30 |
| 1xSH + 1 Cys | 6.20 +/- 0.22 | 1478.20 | 566.30 |
| 2xSH | 8.00 +/- 0.30 | 1479.70 | 566.30 |
| 2xGSH | 5.50 +/- 0.30 | 1522.97 | 566.30 |

WILEY-VCH

| | | | |
|---|---|---|---|
| 1xGSH + 1xCys | 5.50 +/- 0.20 | 1524.78 | 566.30 |
| 1xGSH + 1xSH | 6.30 +/- 0.21 | 1569.80 | 566.30 |

Results are shown on **Table S. 1**.

#### 1.3 Confirmation of the position of the additional interchain disulfide bridge

The redox state of all the cysteines in the Fc-C239i variants was determined by means of non-reducing peptide mapping. Having identified that the inserted cysteine participated in an additional disulfide bond, a targeted LC-MS method was developed to provide improved $MS^2$ fragmentation data to determine the position of the additional disulfide bond.

##### 1.3.1 Non-reduced peptide map preparation: NEM capping, denaturation, and Lys-C digestion

50 µg of iDSB Fc-C239i variants were alkylated using 2.5 µg of N-ethyl maleimide (NEM) at room temperature for 20 minutes. Samples were then dried using a rotary evaporator remove excess NEM. Protein was reconstituted in 45 µL of a denaturing solution (7.1 M GdmCl, 5.6 mM sodium phosphate pH 7.0,0.1 M NaCl) incubated at 37 °C for 30 minutes. The solution was diluted in 125 µL of 100 mM Sodium Phosphate Buffer, pH 7.0 and 0.5 µL of 40 mM EDTA. 2.5 µg of Endoproteinase Lys C were added to the sample and incubated at 37 °C for 2 hours. A further 2.5 µg of Endoproteinase Lys C were added and incubated for a final 2 hours.

##### 1.3.2 LC-MS run

In order to identify the position of the additional disulfide bond observed in the C239i-containing peptide, the non-reduced peptide digest was analysed using a targeted LC-MS method. The digested samples were analysed by means of reverse-phase liquid chromatography (Acquity i-Class UPLC, Waters, Manchester, UK) coupled to mass spectrometry (QExactive Orbitrap ThermoFisher mass spectrometer). Material was separated using Peptide BEH C18 Column, 300 Å 1.7 µm, 2.1 mm x 150 mm over a 12-minute linear gradient of 25-45% B. MS acquisition was performed on a Q-Exactive HFX Mass spectrometer (ThermoFisher, Breman, Germany), over a 200-2000 m/z range, using an AGC target of 3000000 and a maximum injection time of 100 ms. $MS^2$ was acquired in profile mode using an AGC target of 100000 and a maximum injection time of 100 ms. Fragmentation was achieved by HCD using a collision energy of 35%. A targeted $MS^2$ analysis was performed on the 1415.71 (4+) ion, previously identified as the non-reduced 4+ hinge peptide (THTCPPCPAPELLGGPSCVFLFPPKPK-THTCPPCPAPELLGGPSCVFLFPPKPK) containing three disulfide bonds.

The results are represented in **Figure S. 4** and discussed in Supporting Information, section 4.1.

#### 1.4 Quality control of the Fc-C239i enriched formats – state of the inserted cysteine over time in guanidinium chloride

##### 1.4.1 Sample preparation

For each of the three variants, 100 µg were prepared in 34 µL, in 0 or 3.5 M GdmCl, in 20 mM His pH 5.5. They were incubated for 0 or 7 days at 25 °C.

##### 1.4.2 Non-reduced peptide map preparation: NEM capping, denaturation, and Lys-C digestion

After the incubation, the samples were alkylated by adding 5 µg of N-ethylmaleimide (NEM), and incubated at room temperature for 20 min. Samples were then buffer exchanged three times into 7.1 M GdmCl, 5.6 mM phosphate pH 7.0 and 0.1 M NaCl with 10 kDa Amicon filters, and concentrated to a final volume of 90 µL. 250 µL of 100 mM phosphate buffer pH 7.0, 1 µL of 40 mM EDTA and 10 µL of 1 mg/mL of Endoproteinase Lys-C were added to each sample, and incubated for two hours at 37 °C. 10 µL at 1 mg/mL of Lys-C were added again, and incubated for another two hours at 37 °C. After digestion, the material was split in half. 5 µL of 500mM DTT was added to 45µL of digested protein and incubated at room temperature to reduce. 5µL of water was added to 45µL of the same sample to act as a non-reduced control. Both samples were run side by side by LC-MS for comparative analysis.

##### 1.4.3 LC-MS run

The digested samples were analysed by means of reverse-phase liquid chromatography (Acquity i-Class UPLC, Waters, Manchester, UK) coupled to mass spectrometry (Orbitrap Fusion ThermoFisher mass spectrometer). Reduced and non-reduced samples were compared to identify disulfide-containing peptides, and identification was performed using a combination of $MS^1$ and $MS^2$. Peptides were separated using a Peptide BEH C18 Column, 300 Å 1.7 µm, 2.1 mm x 150 mm (Waters, Manchester, UK) over a 76-minute linear gradient of 5-45% B (mobile phase A: 0.02 % TFA in water; mobile phase B: 0.02% TFA in ACN). MS data was acquired over a 250-2000 m/z range, using an AGC target of 200000 and a maximum injection time of 50 ms. $MS^2$ was acquired in the Ion trap in Centroid mode, using an AGC target of 10000 and a maximum injection time of 35 ms. Fragmentation was achieved by CID using a collision energy of 35%. The data was then processed on the Qual Browser Thermo Xcalibur 3.0.63 software.

##### 1.4.4 Mass spectrometry data analysis

The peptides corresponding to the hinge region were searched (NEM-capped, cysteine-capped and disulfide bridged). The presence of the variants was quantitated by combining the most intense isotope on the 4+ m/z and 5+ m/z distribution, and then integrating the area under the obtained peaks on the total ion count chromatogram.

Results are shown in **Figure 1 A** and **Table S. 2**.

#### 1.5 Evolution of the inserted-cysteine states over time upon the denaturant concentrations stress and the incubation time

The proportion of the variants was evaluated by targeted non-reduced peptide mapping.

##### 1.5.1 Sample preparation

The first set of experiments consisted in observing the effect of denaturant concentration after 7 days of incubation at 25 °C on the inserted-cysteine states. For this first set of experiments, 100 µg of the free-thiol enriched and doubly-capped cysteine enriched variants were prepared in 0, 0.5, 1, 1.5, 2, 2.5, 3 and 3.5 M GdmCl, in 20 mM His pH 5.5 in 34 total µL. They were incubated 7 days at 25 °C.

The second set of experiments aimed at assessing the effect of incubation time at 25 °C in 3.5 M GdmCl on the inserted-cysteine states. For the second set of experiments, 100 µg of the free-thiol enriched and doubly-capped cysteine enriched variants were prepared 3.5 M GdmCl, in 20 mM His pH 5.5 in 34 total µL. They were incubated for 0, 1, 2, 3 and 4 days at 25 °C.

##### 1.5.2 Non-reduced peptide map preparation: NEM capping, denaturation and Lys-C digestion

After the incubation, the samples were alkylated adding 5 µg of N-ethylmaleimide (NEM), and incubated at room temperature for 20 min. The samples were buffer exchanged three times into 7.1 M GdmCl, 5.6 mM sodium

WILEY-VCH

phosphate pH 7.0 and 0.1 M NaCl with 10 kDa Amicon filters, and concentrated to a final volume of 60 µL, and incubated at 37°C for 30 minutes. From this volume, 15 µL were taken (25 µg of antibody) and were then diluted by four in 100 mM $NaPO_4$ pH 7.0 containing 0.4% 40 mM EDTA, diluting the guanidinium chloride concentration to 1.8 M in preparation for digest. An aliquot of 10 µL (0.5 µg) of Lys-C was added to ~ 25 µg of thiol capped denatured protein at a 1:50 enzyme:protein ratio (10 and the mixture was incubated at 37 °C for 2 hours. A further 10 µL (0.5 µg) of Lys-C was added and incubated for a final 2 hours. Samples were then analysed.

##### 1.5.3 Reversed Phase LC

Mobile Phase A contained 0.1% TFA in Water and Mobile Phase B contained 0.1% TFA in 100% Acetonitrile. The following LC conditions were used: flow rate 0.15mL/minute, column temperature 55°C throughout the separation with the autosampler maintained at 4°C. Injections of 10 µL of ~ 0.42 mg/mL peptide sample were separated using a UPLC Peptide CSH C18, 130Å pore size, 1.7 µm bead size, 2.1 mm X 150 mm column (Acquity). The gradient started at 0% B until a step up to 24% at 4 minutes and then gradually increased to 26% at 8 minutes, followed by a step up to 80%. The gradient was dropped back to 100% A to equilibrated for the next injection. Total run time per samples was 12 minutes.

##### 1.5.4 Mass spectrometer settings

The Lys-C digested peptides were separated by reversed phase uHPLC coupled to MS (Waters Xevo TQS). The MS desolvation temperature was 600, source cone voltage was 40V, desolvation gas flow 400 L/hr, capillary voltage 3kV, Q3 collision energy 40eV. The different variants were targeted according to the values referenced in the following table.

| Compound | RT (minutes) | *m/z* precursor ion | *m/z* fragment ion |
|---|---|---|---|
| iDSB | 5.70 +/- 0.20 | 1416.50 | 566.30 |
| 2xCys | 5.50 +/- 0.20 | 1476.70 | 566.30 |
| 1xSH + 1 Cys | 6.20 +/- 0.22 | 1478.20 | 566.30 |
| 2xSH | 8.00 +/- 0.30 | 1479.70 | 566.30 |
| 2xGSH | 5.50 +/- 0.30 | 1522.97 | 566.30 |
| 1xGSH + 1xCys | 5.50 +/- 0.20 | 1524.78 | 566.30 |
| 1xGSH + 1xSH | 6.30 +/- 0.21 | 1569.80 | 566.30 |

Results are shown in **Figure 1 B** and **Figure S. 5**.

#### 1.6 Measurement of the thermodynamic stability of the antibodies: chemical unfolding and refolding curves

##### 1.6.1 Methods

Chemical unfolding and refolding curves in guanidinium chloride were performed in duplicate or triplicate. Each experiment was composed of forty-one points (120 µL total volume) of increasing concentrations of GdmCl from 0 to 3.5 or 4 M final concentration. 110 µL of denaturant solution was mixed with 10 µL of protein to a final concentration of 1 µM. For the unfolding curves, the stock protein solution was made in 20 mM histidine buffer pH 5.5; for the refolding curves, the protein was first denatured in 5 M GdmCl for fifteen minutes at room temperature and then dispensed into the same forty-one-point denaturant solutions.

The solutions were dispensed with a liquid handling robot (Microlab®500 Series, ML541C, Hamilton Company).

WILEY-VCH

The denaturant solutions mixed with the protein were incubated at 25 °C at different time points until they reached equilibrium (7 days). Each of the forty-one denaturation points were measured in a 100 µL quartz cuvette (Hellma, Precision Cell in Quartz SUPRASIL®, Typ No: 105.250-QS, Light Path: 10x2 mm, Centre: 20 mm). The fluorescence was recorded with a Cary 400 Eclipse Fluorescence Spectrophotometer (Agilent Technologies) thermostatted at 25 °C controlled by a heat block. The samples were excited at 280 nm, the emission was recorded from 300 to 400 nm, with a scan rate of 300 nm $min^{-1}$, excitation and emission band passes were set at 10 nm.

##### 1.6.2 Data Analysis

When the protein is denatured with increasing concentrations of chemical denaturant, the maximum of the fluorescence signal shifts towards red wavelengths. In the native state, the wavelength of maximum fluorescence intensity is at approximately 335 nm and in the denatured state, the wavelength of maximum fluorescence is around 360 nm.  The data were analyzed using an average emission wavelength (AEW), which is the arithmetic mean of the wavelengths weighted by the fluorescence intensity at each wavelength. It is calculated as shown in Equation 1:

$$AEW = \frac{\sum_{i=1}^{N} F_i.\lambda_i}{\sum_{i=1}^{N} F_i} \qquad \textbf{(1)}$$

where $F_i$ is the intensity of fluorescence at the wavelength $i$, and $\lambda_i$ the wavelength.

A three-state model in which an intermediate state between the native and denatured state is sufficiently stable to be populated and observed was used. The equilibrium between the different species is as follows:

$$\mathrm{N} \overset{K_{I-N}}{\rightleftharpoons} \mathrm{I} \overset{K_{D-I}}{\rightleftharpoons} \mathrm{D} \qquad \textbf{(2)}$$

The denaturation curves, using the AEW data, were fitted to a three-state model using Equation 3[30,31]:

$$F = \frac{\alpha_N+\beta_N.[den]+\alpha_I.\exp\left(\frac{m_{I-N}}{RT}.([den]-[den]_{50\%\,I-N})\right)+(\alpha_D+\beta_D.[den])\ \exp\left(\frac{m_{I-N}}{RT}.([den]-[den]_{50\%\,I-N})\right)\exp\left(\frac{m_{D-I}}{RT}.([den]-[den]_{50\%\,D-I})\right)}{1+\exp\left(\frac{m_{I-N}}{RT}.([den]-[den]_{50\%\,I-N})\right)+\exp\left(\frac{m_{I-N}}{RT}.([den]-[den]_{50\%\,I-N})\right)\exp\left(\frac{m_{D-I}}{RT}.([den]-[den]_{50\%\,D-I})\right)} \qquad \textbf{(3)}$$

where $\alpha_N, \alpha_I, \alpha_D$ are the fluorescence of the native, intermediate and denatured states in $H_2O$ respectively; $\beta_N, \beta_D$ are the slopes of native, intermediate and denatured baselines respectively; $m_{I-N}, m_{D-I}$ are the $m$-values between the intermediate and native state, and denatured and intermediate states respectively; $\Delta G_{I-N}^{H2O}, \Delta G_{D-I}^{H2O}, \Delta G_{D-N}^{H2O}$ are the differences in Gibbs free energy between the intermediate and native states, denatured and intermediate states, and denatured and native states, respectively; $T$ the temperature and $R$ the gas constant.

#### 1.7 Measurement of the thermal stability of the antibodies: differential scanning calorimetry

##### 1.7.1 Methods

The thermal denaturation of NIST mAb Fc, 2xSH Fc-C239i, 2xCys Fc-C239i, iDSB Fc-C239i, as well as 2xSH capped by N-ethylmaleimide (NEM) to avoid interconversion of the cysteine state during the thermal denaturation, were monitored by differential scanning calorimetry (DSC) in triplicates (duplicates for 2xSH NEM capped Fc-C239i), with a Malvern MicroCal VP-DSC instrument. 500 µL of protein at 0.5-5 mg mL$^{-1}$ in 20 mM histidine pH 5.5 were used for each run. For each protein, the baseline was measured first, which consists of buffer in both cells (buffer *versus* buffer), and then the protein was run (buffer *versus* protein). Several clean-up cycles with water and suitability controls with lysozyme at 3 mg mL$^{-1}$ in water were employed before, and after, the actual experiment with the antibody. Each protein sample was scanned twice to investigate the thermal reversibility. The temperature was ramped from 25 to 100 °C, increasing by 95 °C h$^{-1}$. The pre-scan thermostat was set to 2 min, no post-scan thermostat was employed. All the different protein constructs were run in triplicate. The thermal unfolding is not reversible, as the trace of the reheated sample did not overlap with the initial one (Figure S. 8).

##### 1.7.2 Data Analysis

The data were processed with the Origin version 7.0 SR4 software. The baseline thermogram (buffer *versus* buffer) was subtracted from the thermogram of the protein (buffer *versus* protein). Two baselines, one at the beginning and one at the end of the thermogram, were placed to adjust the data, which was then normalized using the concentration of the protein. The unfolding peaks were selected and the thermogram was fitted to the "Non 2-state" model to obtain the melting temperatures ($T_m$) and the enthalpy of unfolding at the $T_m$, $\Delta H_{cal}$.

#### 1.8 Measurement of the kinetic stability of the antibodies: unfolding kinetics

##### 1.8.1 Methods

For the stopped-flow experiments, the native protein and the denaturant solutions were mixed in a 1:10 ratio respectively. Seven stock solutions of guanidinium chloride (GdmCl) were prepared in 20 mM histidine buffer pH 5.5 so that the final concentrations range from 5.5 to 7.0 M GdmCl with an interval of 0.25 M. The stock solutions are 1.1-fold more concentrated than the final solutions, as solutions were diluted by a factor of 10/11 in the rapid mixing step. The protein stock solutions were prepared between 7 and 11 µM to achieve a final concentration between 0.5 to 1 µM after mixing with the denaturant solutions.

The unfolding kinetics were monitored with a SX20 stopped-flow spectrometer from Applied Photophysics (software: SX Spectrometer Control Panel Application version 2.2.27). The temperature of the water bath was set to 25 °C, the excitation wavelength was set to 280 nm, both slit widths were 2 mm. A cut off filter of 320 nm was used. Three short time traces were recorded with pressure hold (2 seconds), to accurately measure the fastest unfolding phase, and three longer time traces (120 sec) were acquired to accurately measure the slower unfolding phase, both at each denaturant concentration.

##### 1.8.2 Fitting of unfolding kinetics

The unfolding curves were fitted with the software Pro DataViewer version 4.2.27. The fluorescence signal corresponding to the unfolding was fitted with a double-exponential function (Equation 4).

$$A(t) = A_1 \,.\exp(-k_1 t) + A_2 \,.\exp(-k_2 t) + c \quad \textbf{(4)}$$

where $A_1$ and $A_2$ are the amplitudes, $k_1$ and $k_2$ the respective unfolding rate constants and $c$ the offset.

The natural logarithm of the rate constants was then calculated and plotted *versus* the corresponding denaturant concentration, and the data fitted with Equation 5[32].

$$\ln k_U^{[den]} = \ln k_U^{H_2O} + m_{k_U}[den] \quad \textbf{(5)}$$

Where $k_U^{[den]}$ is the observed unfolding rate constant at the denaturant concentration $[den]$, $k_U^{H_2O}$ is the unfolding rate constant in water and $m_{k_U}$ is the slope of the plot of ln$k_U^{[den]}$ *versus* denaturant concentration.

#### 1.9 Fast Hydrogen-Deuterium Exchange Mass Spectrometry

The peptide map was done on the 2xCys enriched Fc-239iC variant, with equilibration buffer (20 mM His pH 5.5 in $H_2O$), using a data dependent acquisition (DDA) $MS^2$ approach. MS data was acquired in the Orbitrap Fusion (ThermoFisher) over a 300-2000 m/z range, using an AGC target of 200000 and a maximum injection time of 100 ms. MS2 was acquired in the Ion trap in Centroid mode, using an AGC target of 10000 and and a maximum injection time of 35 ms. Fragmentation was achieved by HCD using a collision energy of 30%. The labelled data on the three enriched variants 2xSH, 2xCys, iDSB Fc-C239i and the wild-type NIST mAb Fc was recorded after 1000, 6000, 30000, 60000, 300000, 600000 and 900000 ms of incubation at 20 °C in deuterated buffer (in 20 mM His pD 5.5 in $D_2O$) with a $MS^1$ method (300-2000 m/z range, AGC target of 200000 and maximum injection time of 100 ms) to avoid scrambling of the deuterations. All data points were run in triplicates.

On the ms2min HDX system[21], patent (WO2020074863A1), 10 µL of protein at 5 µM in equilibration buffer (20 mM His pH 5.5 in $H_2O$) were diluted 20-fold at 20 °C into equilibration or labelling buffer (20 mM His pH 5.1 (pD 5.5) in $D_2O$), to generate peptide map or labelled data respectively. The mixture was then diluted 1:1 with quench buffer at 2 °C (100 mM His, 8 M urea, 0.5 M TCEP, pH 2.5 when mixed 1:1 with labelling buffer at 4 °C). The quench solution was then injected into Waters nanoAcquity UPLC system, flowing for 4 min at 50 µL/min onto the pepsin column at 20 °C for digestion (Waters Enzymate™ BEH Pepsin Column (2.1 x 30 mm, 5 µm)) to the trap (pushed by LC-MS grade $H_2O$ with 0.2% formic acid, and then eluting from the C18 analytical column for 10 minutes, from a 5% to 40 % organic phase (ACN with 0.2% formic acid).

The peptide map was generated on BioPharmaFinder from equilibration data (in 20 mM His pH 5.5 in $H_2O$) acquired with a $MS^2$ method. The peptides obtained where filtered by the MS identification method ($MS^2$ only) and the peptide mass error lower than 10 ppm. The exported csv with the peptides as well as that same undeuterated data were imported into HDExaminer, to operate a second filtration of the peptides: only the charge state with the highest intensity from the peptide map data was kept per peptide for comparative accuracy between the charge states, selected according the highest intensity, the sharpest extracted ion chromatogram. After the peptide pool was curated, the labelled data was added, and the D incorporation per peptide data was then exported as a csv file. Given that the N-terminus of the Fc domain was different for the wild-type NIST mAb Fc and the Fc-C239i variants, the first peptides don't exist for the wild-type and the data comparison starts at residue Lys 242. We employed two complementary methods of analysis to identify which deuterium incorporations were significant, to observe the deuterium exchange for each time point separately and overall.

The first method used, which results are represented and discussed in the main body of the paper (Figure 3) was first described by Dobson, Devine, Phillips *et al.*, 2016[33]. Starting with a csv file containing the D incorporation data, this Matlab-coded method first assesses if the incorporation of deuterium per peptide is significant compared to the wild-type (assessed by a t-test, if p-value < 0.05), then sums the significant time points per peptide, subsequently converts the peptide D incorporation to amino acid incorporation by dividing the D incorporation by the maximal number of D that can be exchanged per peptide, subtracting the D incorporation from the wild-type and dividing the amino acid incorporation by the redundancy and finally normalizing it. The output is the deuterium incorporation for each individual amino acid. The crystal structure of the Fc domain (PDB:3AVE) was coloured with the red-white-blue scale according to the relative incorporation of deuterium per

WILEY-VCH

amino. The additional cysteine was removed from the incorporation data to map on the structure. The differential plots obtained are presented in **Figure S. 11**, and the summary HDX-MS parameters are in **Table S. 10**.

The second method used was first described in Cornwell, O., et al., 2018[34], and represents the deuterium incorporation per timepoint, separately, rather than summing the incorporations. From the csv file containing the D incorporation data per peptide, this approach calculates the combined mean relative fractional uptake (deuterium uptake corrected for the number of exchangeable amides in the peptide) for each amino acid, by averaging the relative fractional uptake values for peptides which cover the amino acid in question. A combined standard deviation is then calculated from the standard deviations for each of the peptides from replicate measurements, and the variances between charge states from the same measurement. This process is repeated for each residue in the sequence, for every measured time point, and all states. To identify the statistically significant differences for each amino acid, per time point and between the states and the wild-type (reference), statistical analysis can be performed using one way ANOVA and post hoc Tukey tests ($p$-value < 0.05), using the combined mean relative fractional uptake, combined standard deviation and total number of measurements for each residue (n = replicates x peptides covering the residue). Only the significant D incorporation differences (between the variants and WT) are plotted onto the Fc domain crystal structure (PDB:3AVE) with the red-white-blue scale, for each variant and each timepoint. The additional cysteine was removed from the incorporation data to map on the structure. The results with this method are in **Figure S. 13**.

WILEY-VCH

#### 2 Supplementary Figures

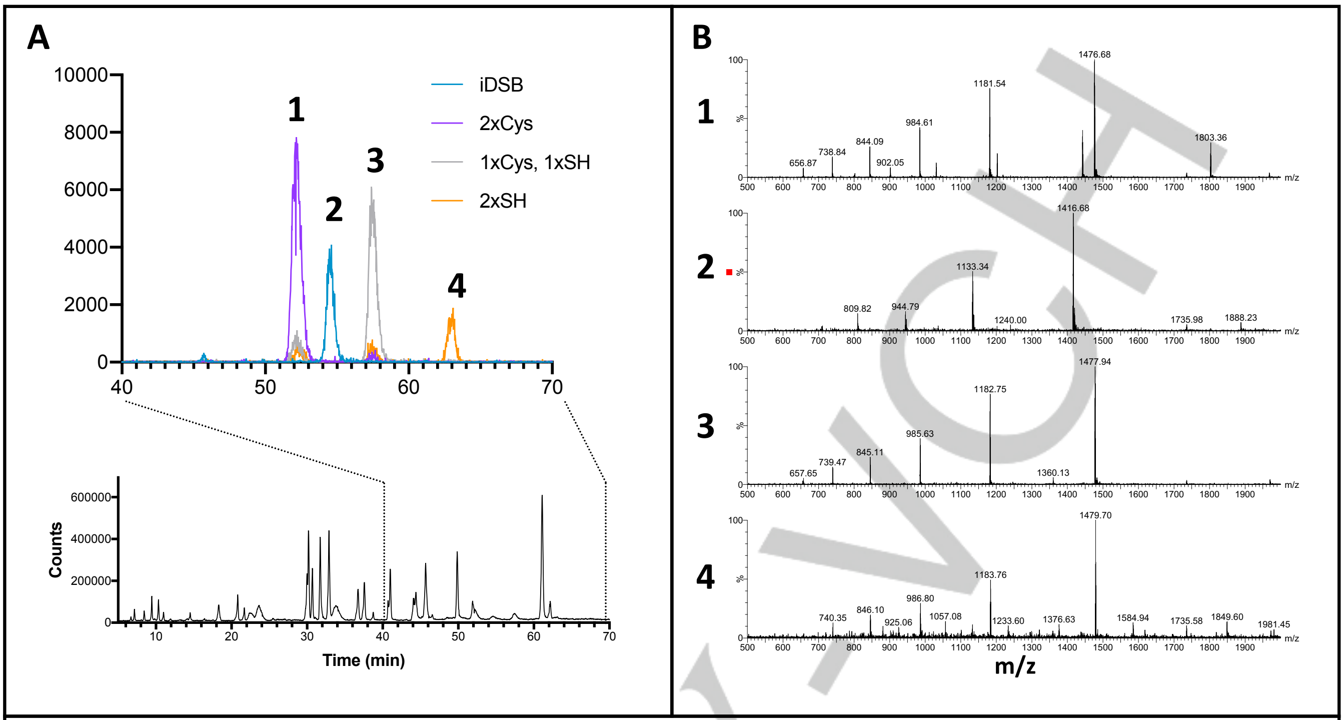


C

| Peptide diagnostic of thiol state | Theoretical monoisotopic mass (Da) | Theoretical average mass of 4+ ion |
|---|---|---|
| [THTCPPCPAPELLGGPSC(Cys)VFLFPPKPK]=[THTCPPCPAPELLGGPSC(Cys)VFLFPPKPK] | 5899.09 | 1476.76 |
| [THTCPPCPAPELLGGPSC(iDSB)VFLFPPKPK]=[THTCPPCPAPELLGGPSC(iDSB)VFLFPPKPK] | 5658.78 | 1416.69 |
| [THTCPPCPAPELLGGPSC(Cys)VFLFPPKPK]=[THTCPPCPAPELLGGPSC(SH*)VFLFPPKPK] | 5905.07 | 1478.26 |
| [THTCPPCPAPELLGGPSC(SH*)VFLFPPKPK]=[THTCPPCPAPELLGGPSC(SH*)VFLFPPKPK] | 5911.05 | 1479.75 |

**Figure S. 1:** Non-reduced peptide mapping of Fc-C239i antibody. **A.** Total ion chromatogram (bottom) and zoom of extracted ion chromatograms (top) for iDSB, 2xCys, 1xCys & 1xSH and 2xSH thiol states (1 to 4). **B.** Mass spectrum for each extracted ion (1 to 4). **C.** Theoreticasl MW for peptide diagnostic of thiol state; * denotes modification by N-Ethylmaleimide.

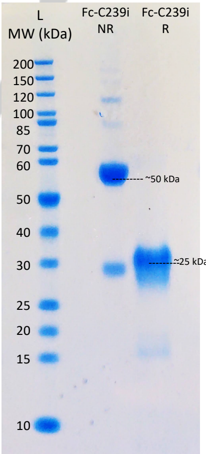


**Figure S. 2:** SDS-Page gel of 2xCys Fc-C239i. NR: non reduced. R: reduced. L: ladder: Page Ruler™ unstained ladder, Thermo Scientific.

WILEY-VCH

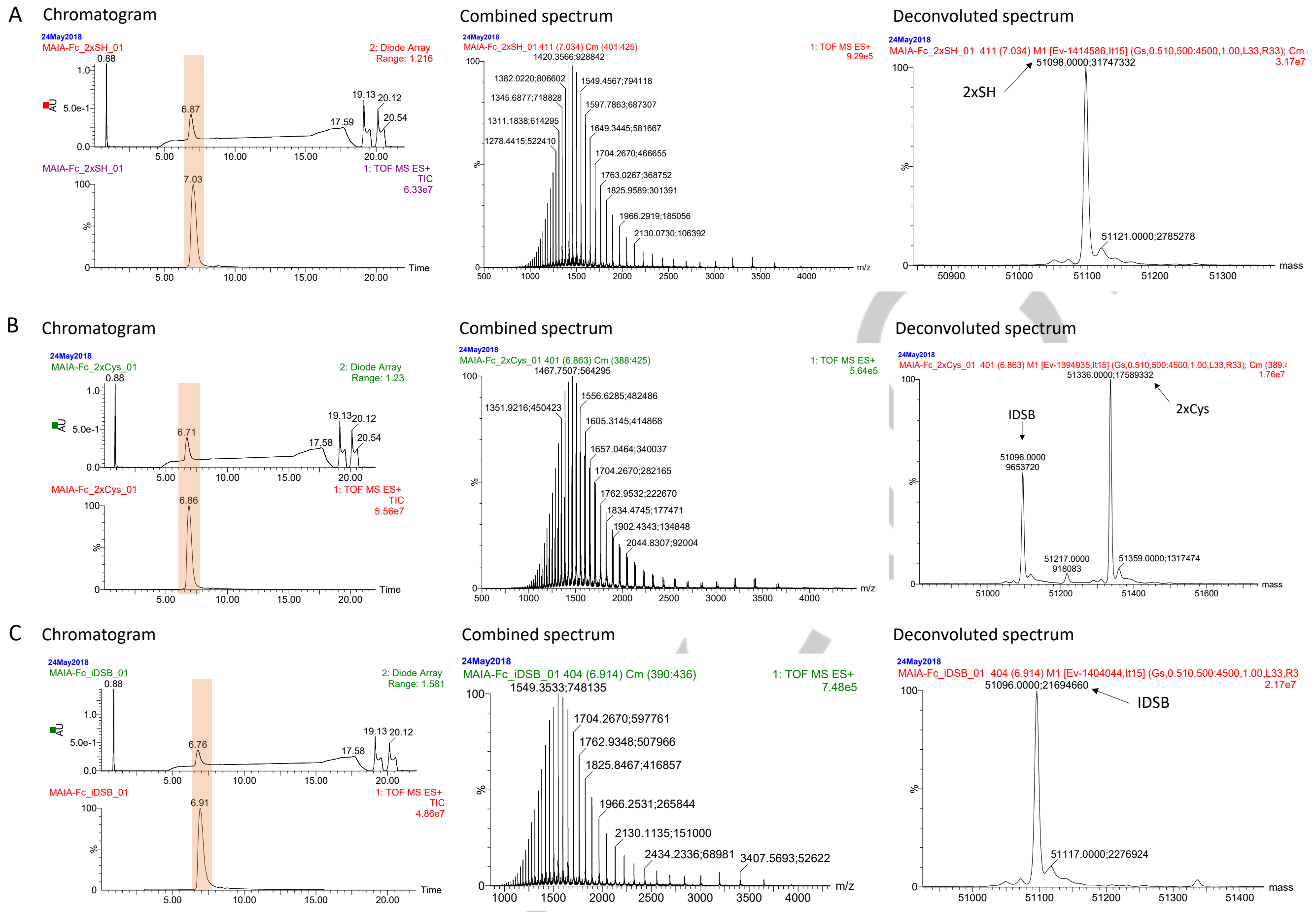


**Figure S. 3:** Chromatograms, combined spectra and deconvoluted spectra (LC-MS) of A. 2xSH Fc-C239i; B. 2xCys Fc-C239i; C. iDSB Fc-C239i enriched variants. The highlighted peak on the chromatogram is the portion that was combined.

WILEY

**A**

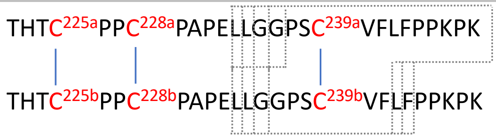


**B**

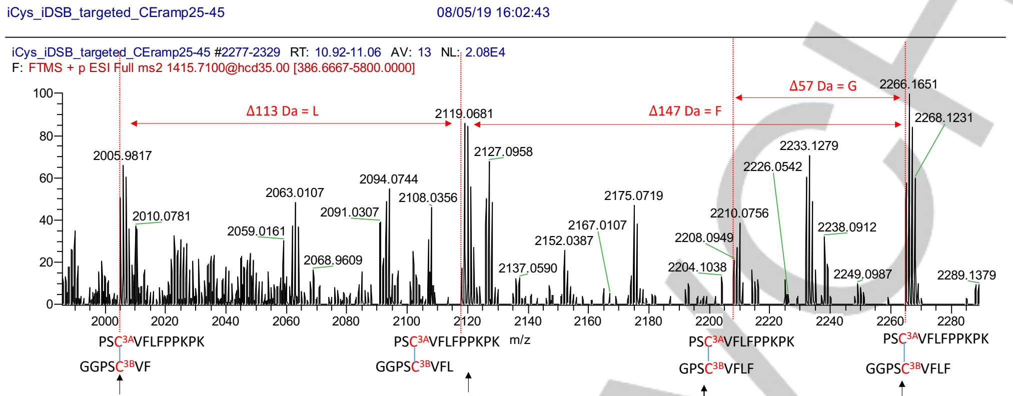


| Peptides | Charge state | Oxidised SS | Th. Sum | Exp. | ppm |
|---|---|---|---|---|---|
| PSCVFLFPPKPK | neutral | 1357.73 | 2005.004 | 2004.9803 | -11.8 |
| GGPSCVF | b7 +1 | 647.274 | | | |

| Peptides | Charge state | Oxidised SS | Th. Sum | Exp. | ppm |
|---|---|---|---|---|---|
| PSCVFLFPPKPK | neutral | 1357.73 | 2118.088 | 2118.044 | -20.8 |
| GGPSCVFL | b8 +1 | 760.358 | | | |

| Peptides | Charge state | Oxidised SS | Th. Sum | Exp. | ppm |
|---|---|---|---|---|---|
| PSCVFLFPPKPK | neutral | 1357.73 | 2265.156 | 2265.1251 | -13.6 |
| GGPSCVFLF | b ion +1 | 907.426 | | | |

| Peptides | Charge state | Oxidised SS | Th. Sum | Exp. | ppm |
|---|---|---|---|---|---|
| PSCVFLFPPKPK | neutral | 1357.73 | 2208.135 | 2208.0949 | -18.2 |
| GPSCVFLF | b ion +1 | 850.405 | | | |

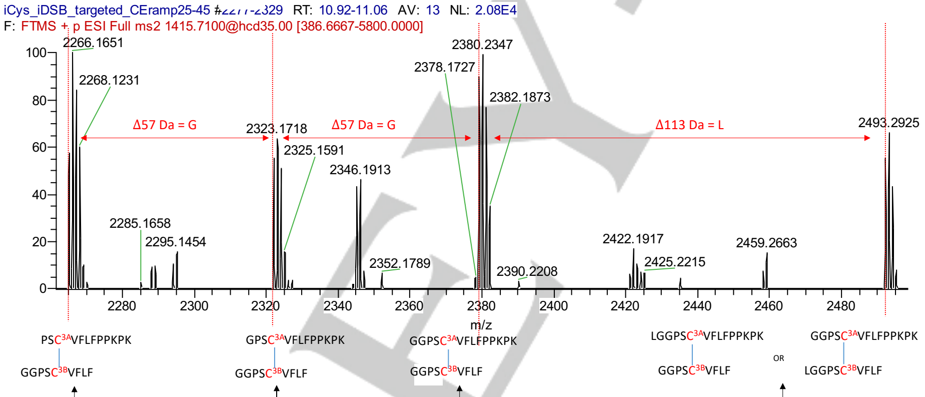


| Peptides | Charge state | Oxidised SS | Th. Sum | Exp. | ppm |
|---|---|---|---|---|---|
| PSCVFLFPPKPK | neutral | 1357.73 | 2265.156 | 2265.1251 | -13.6 |
| GGPSCVFLF | b ion +1 | 907.426 | | | |

| Peptides | Charge state | Oxidised SS | Th. Sum | Exp. | ppm |
|---|---|---|---|---|---|
| GGPSCVFLFPPKPK | neutral | 1471.78 | 2379.206 | 2379.2357 | 12.5 |
| GGPSCVFLF | b ion +1 | 907.426 | | | |

| Peptides | Charge state | Oxidised SS | Th. Sum | Exp. | ppm |
|---|---|---|---|---|---|
| LGGPSCVFLFPPKPK | neutral | 1584.86 | 2492.286 | 2492.2735 | -5.0 |
| GGPSCVFLF | b ion +1 | 907.426 | | | |
| GGPSCVFLFPPKPK | neutral | 1471.78 | 2492.29 | 2492.2735 | -6.6 |
| LGGPSCVFLF | b ion +1 | 1020.51 | | | |

| Peptides | Charge state | Oxidised SS | Th. Sum | Exp. | ppm |
|---|---|---|---|---|---|
| GPSCVFLFPPKPK | neutral | 1414.76 | 2322.186 | 2322.1414 | -19.9 |
| GGPSCVFLF | b ion +1 | 907.426 | | | |

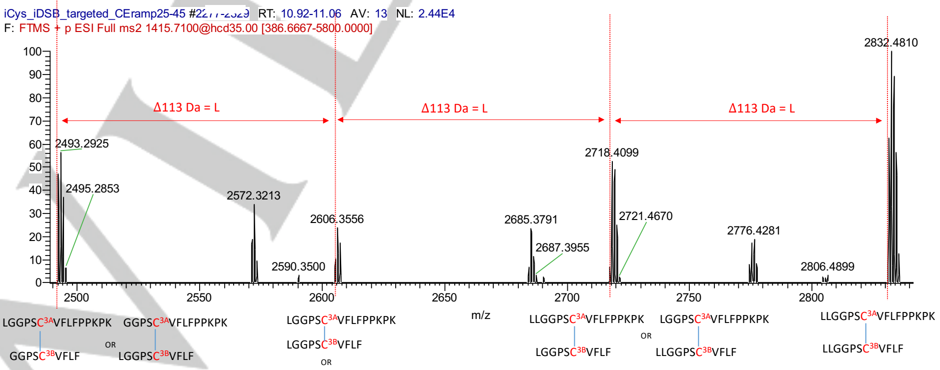


| Peptides | Charge state | Oxidised SS | Th. Sum | Exp. | ppm |
|---|---|---|---|---|---|
| LGGPSCVFLFPPKPK | neutral | 1584.86 | 2492.286 | 2492.2735 | -5.0 |
| GGPSCVFLF | b ion +1 | 907.426 | | | |
| GGPSCVFLFPPKPK | neutral | 1471.78 | 2492.29 | 2492.2735 | -6.6 |
| LGGPSCVFLF | b ion +1 | 1020.51 | | | |

| Peptides | Charge state | Oxidised SS | Th. Sum | Exp. | ppm |
|---|---|---|---|---|---|
| LGGPSCVFLFPPKPK | neutral | 1584.86 | 2605.37 | 2605.3455 | -9.4 |
| LGGPSCVFLF | b ion +1 | 1020.51 | | | |
| LLGGPSCVFLFPPKPK | neutral | 1697.95 | 2605.376 | 2605.3455 | -11.7 |
| GGPSCVFLF | b ion +1 | 907.426 | | | |
| GGPSCVFLFPPKPK | neutral | 1471.78 | 2605.374 | 2605.3455 | -10.9 |
| LLGGPSCVFLF | b ion +1 | 1133.594 | | | |

| Peptides | Charge state | Oxidised SS | Th. Sum | Exp. | ppm |
|---|---|---|---|---|---|
| LLGGPSCVFLFPPKPK | neutral | 1697.95 | 2718.46 | 2718.4099 | -18.4 |
| LGGPSCVFLF | 'b10' +1 | 1020.51 | | | |
| LGGPSCVFLFPPKPK | neutral | 1584.86 | 2718.454 | 2718.4099 | -16.2 |
| LLGGPSCVFLF | 'b11' +1 | 1133.594 | | | |

| Peptides | Charge state | Oxidised SS | Th. Sum | Exp. | ppm |
|---|---|---|---|---|---|
| LLGGPSCVFLFPPKPK | neutral | 1697.95 | 2831.544 | 2831.4828 | -21.6 |
| LLGGPSCVFLF | 'b11' +1 | 1133.594 | | | |

9

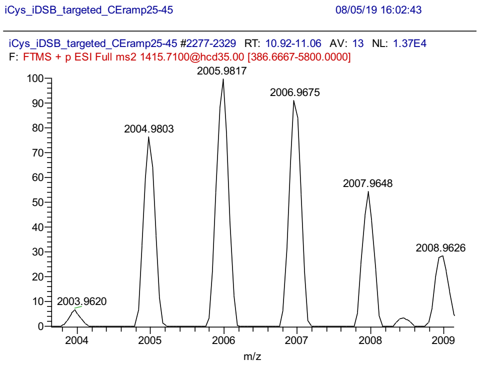


PS$C^{3A}$VFLFPPKPK

GGPS$C^{3B}$VF

| Peptides | Charge state | Oxidised SS | Sum | Experimental | ppm |
|---|---|---|---|---|---|
| PSCVFLFPPKPK | neutral | 1357.73 | 2005.004 | 2004.9803 | -11.8 |
| GGPSCVF | b ion +1 | 647.274 | | | |

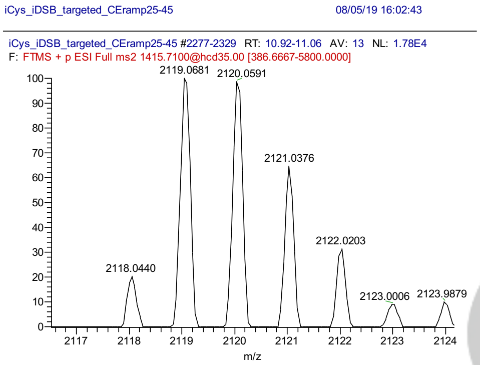


PS$C^{3A}$VFLFPPKPK

GGPS$C^{3B}$VFL

| Peptides | Charge state | Oxidised SS | Sum | Experimental | ppm |
|---|---|---|---|---|---|
| PSCVFLFPPKPK | neutral | 1357.73 | 2118.088 | 2118.044 | -20.8 |
| GGPSCVFL | b ion +1 | 760.358 | | | |

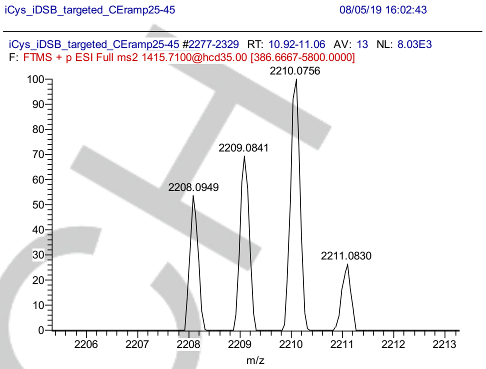


PS$C^{3A}$VFLFPPKPK

GPS$C^{3B}$VFLF

| Peptides | Charge state | Oxidised SS | Sum | Experimental | ppm |
|---|---|---|---|---|---|
| PSCVFLFPPKPK | neutral | 1357.73 | 2208.135 | 2208.0949 | -18.2 |
| GPSCVFLF | b ion +1 | 850.405 | | | |

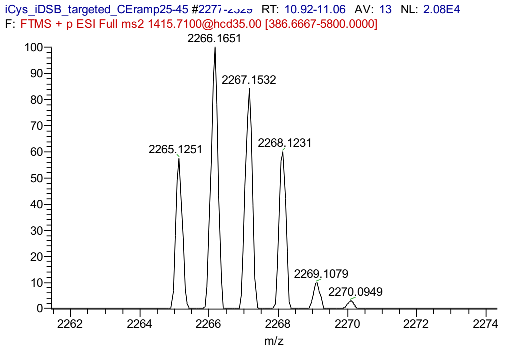


PS$C^{3A}$VFLFPPKPK

GGPS$C^{3B}$VFLF

| Peptides | Charge state | Oxidised SS | Sum | Experimental | ppm |
|---|---|---|---|---|---|
| PSCVFLFPPKPK | neutral | 1357.73 | 2265.156 | 2265.1251 | -13.6 |
| GGPSCVFLF | b ion +1 | 907.426 | | | |

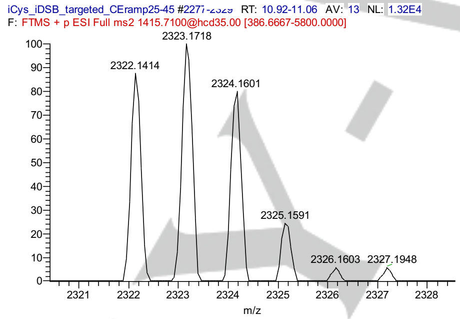


GPS$C^{3A}$VFLFPPKPK

GGPS$C^{3B}$VFLF

| Peptides | Charge state | Oxidised SS | Sum | Experimental | ppm |
|---|---|---|---|---|---|
| GPSCVFLFPPKPK | neutral | 1414.76 | 2322.186 | 2322.1414 | -19.2 |
| GGPSCVFLF | b ion +1 | 907.426 | | | |

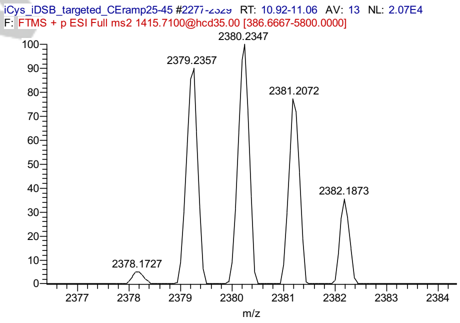


GGPS$C^{3A}$VFLFPPKPK

GGPS$C^{3B}$VFLF

| Peptides | Charge state | Oxidised SS | Sum | Experimental | ppm |
|---|---|---|---|---|---|
| GGPSCVFLFPPKPK | neutral | 1471.78 | 2379.206 | 2379.2357 | 12.5 |
| GGPSCVFLF | b ion +1 | 907.426 | | | |

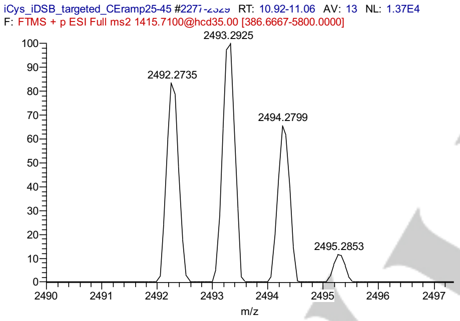


LGGPS$C^{3A}$VFLFPPKPK / GGPS$C^{3B}$VFLF

OR

GGPS$C^{3A}$VFLFPPKPK / LGGPS$C^{3B}$VFLF

| Peptides | Charge state | Oxidised SS | Sum | Experimental | ppm |
|---|---|---|---|---|---|
| LGGPSCVFLFPPKPK | neutral | 1584.86 | 2492.286 | 2492.2735 | -5.0 |
| GGPSCVFLF | b ion +1 | 907.426 | | | |
| GGPSCVFLFPPKPK | neutral | 1471.78 | 2492.29 | 2492.2735 | -6.6 |
| LGGPSCVFLF | b ion +1 | 1020.51 | | | |

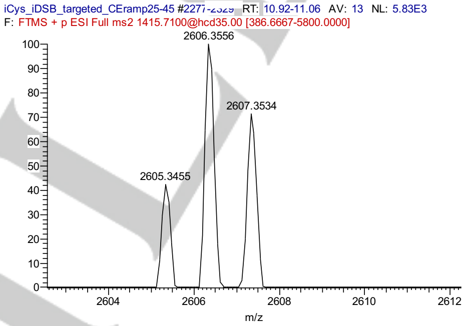


LGGPS$C^{3A}$VFLFPPKPK / LGGPS$C^{3B}$VFLF

OR

LLGGPS$C^{3A}$VFLFPPKPK / GGPS$C^{3B}$VFLF

OR

GGPS$C^{3A}$VFLFPPKPK / LLGGPS$C^{3B}$VFLF

| Peptides | Charge state | Oxidised SS | Sum | Experimental | ppm |
|---|---|---|---|---|---|
| LGGPSCVFLFPPKPK | neutral | 1584.86 | 2605.37 | 2605.3455 | -9.4 |
| **LGGPSCVFLF** | **b ion +1** | **1020.51** | | | |
| LLGGPSCVFLFPPKPK | neutral | 1697.95 | 2605.376 | 2605.3455 | -11.7 |
| **GGPSCVFLF** | **b ion +1** | **907.426** | | | |
| GGPSCVFLFPPKPK | neutral | 1471.78 | 2605.374 | 2605.3455 | -10.9 |
| **LLGGPSCVFLF** | **b ion +1** | **1133.594** | | | |

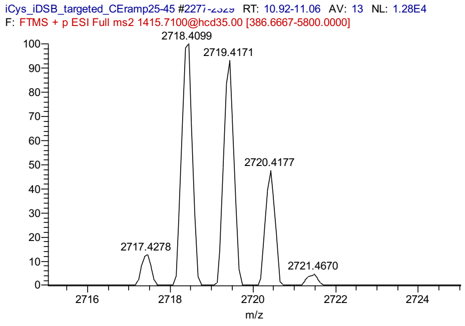


LLGGPS$C^{3A}$VFLFPPKPK / LGGPS$C^{3B}$VFLF

OR

LGGPS$C^{3A}$VFLFPPKPK / LLGGPS$C^{3B}$VFLF

| Peptides | Charge state | Oxidised SS | Sum | Experimental | ppm |
|---|---|---|---|---|---|
| LLGGPSCVFLFPPKPK | neutral | 1697.95 | 2718.46 | 2718.4099 | -18.4 |
| LGGPSCVFLF | 'b10' +1 | 1020.51 | | | |
| LGGPSCVFLFPPKPK | neutral | 1584.86 | 2718.454 | 2718.4099 | -16.2 |
| LLGGPSCVFLF | 'b11' +1 | 1133.594 | | | |

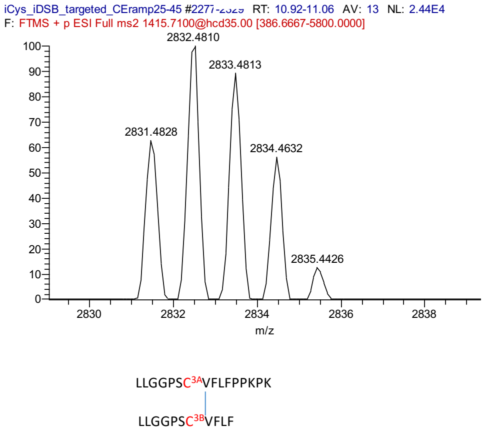


| Peptides | Charge state | Oxidised SS | Sum | Experimental | ppm |
|---|---|---|---|---|---|
| LLGGPSCVFLFPPKPK | neutral | 1697.95 | 2831.544 | 2831.4828 | -21.6 |
| LLGGPSCVFLF | 'b11' +1 | 1133.594 | | | |

**Figure S. 4:** Identification of iDSB by tandem mass spectrometry. **A.** Sequence of the hinge peptide fragmented by tandem mass spectrometry, with the fragmented ions identified. **B.** Reconstitution of the sequence fragmented by $MS^2$ amino by amino acid. **C.** Detailed m/z spectra for each main peak.

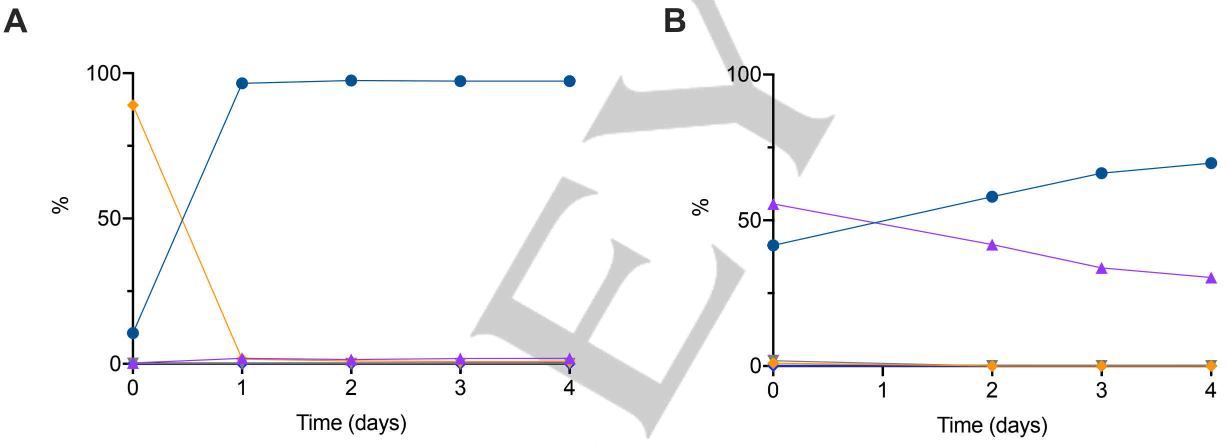


**Figure S. 5:** Effect of the incubation time at 25 °C in 3.5 M GdmCl on (A) 2xSH Fc-C239i enriched starting material and (B) 2xCys Fc-C239i enriched starting material. Blue circle: iDSB Fc-C239i; purple triangle: 2xCys Fc-C239i; orange lozenge: 2xSH Fc-C239i; grey triangle: single free-thiol Fc-C239i. The doubly-glutathione capped Fc-C239i, one glutathione and one cysteine Fc-C239i and single free-thiol and capped by one glutathione Fc-C239i forms were monitored but not detected.

WILEY-VCH

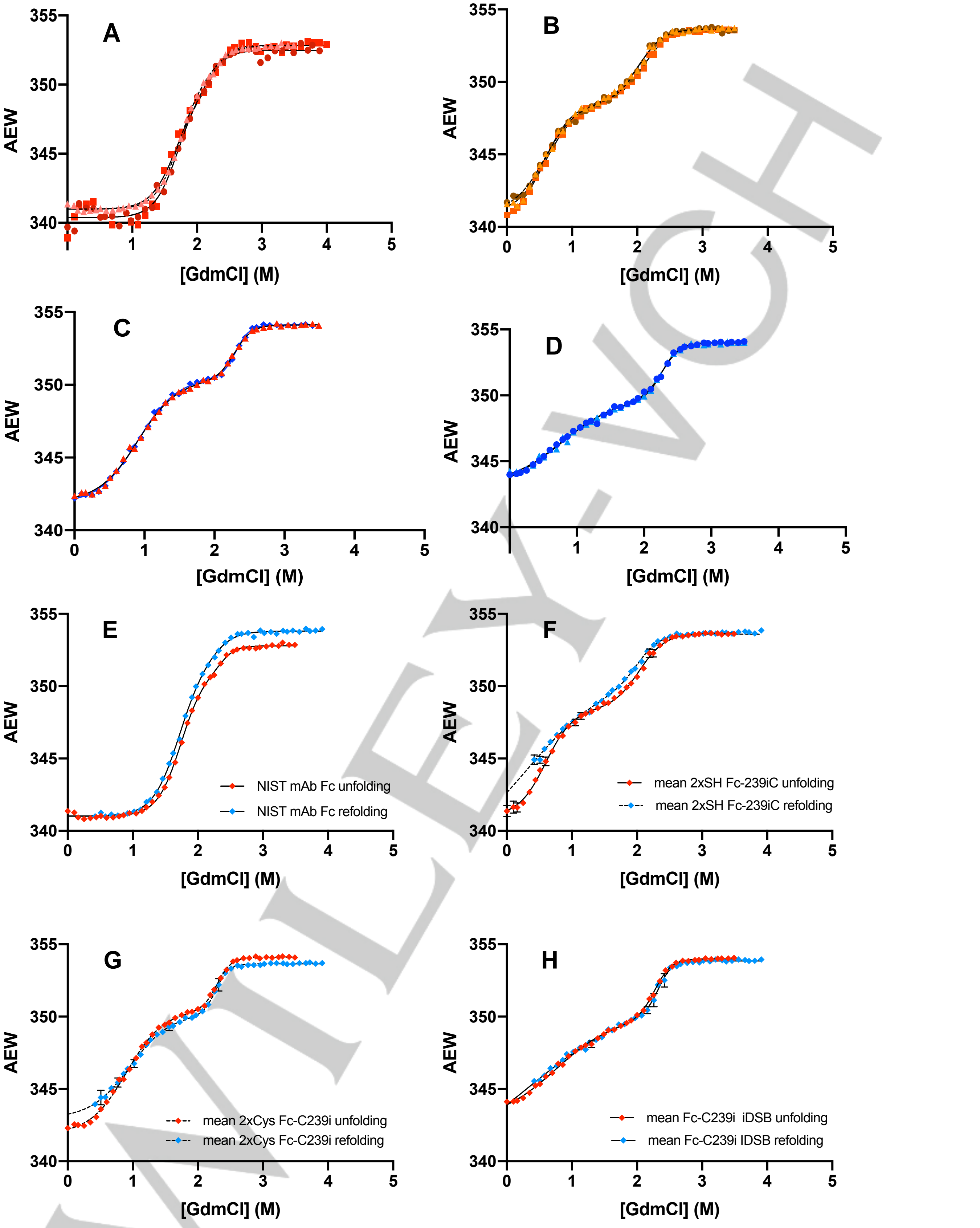


WILEY-VCH

**Figure S. 6:** Results of the chemical denaturation curves. All the denaturation curves were done at 1 µM final protein concentration in 20 mM His pH 5.5 and samples were incubated at 25 °C for seven days. **A.** Triplicates of the unfolding curves of NIST mAb Fc (WT) after seven days of incubation. **B.** Triplicates of the unfolding curves of

2xSH Fc-C239i enriched variant after seven days of incubation. **C.** Duplicates of the unfolding curves of 2xCys Fc-C239i enriched variant after seven days of incubation. **C.** Duplicates of the unfolding curves of iDSB Fc-C239i enriched variant after seven days of incubation. **E.** Unfolding (red – one curve) and refolding (blue – one curve) curves of NIST mAb Fc (WT) after 7 days of incubation. **F.** Unfolding (red - mean of triplicates) and refolding (blue - mean of triplicates) curves of 2xSH Fc-C239i. **G.** Unfolding (red – one curve) and refolding (blue - mean of triplicates) curves of 2xCys Fc-C239i. **H.** Unfolding (red – mean of duplicates) and refolding (blue - mean of duplicates) curves of iDSB Fc-C239i.

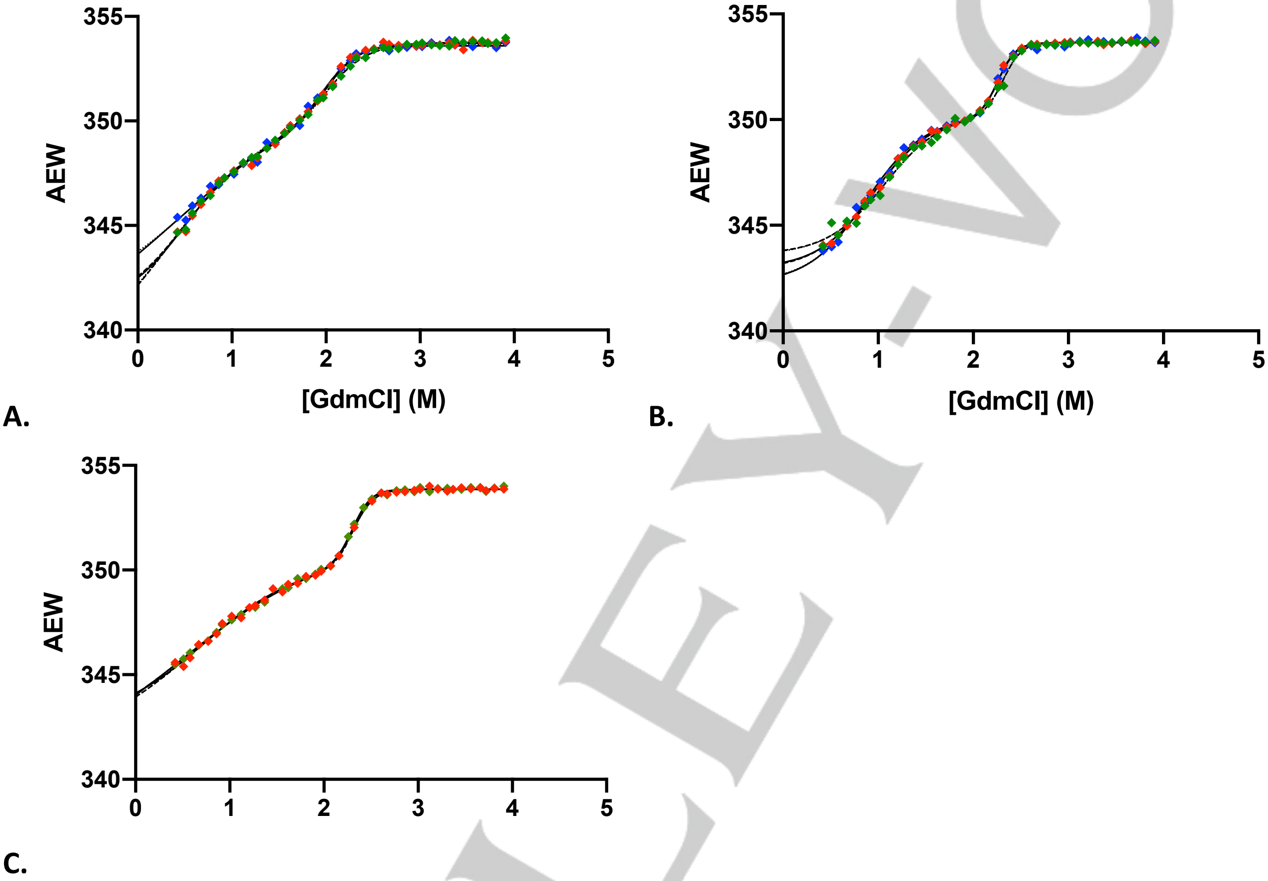


**Figure S. 7:** Repeats of chemical renaturation curves of the Fc-C239i formats. All these denaturation curves were done at 1 µM final of protein in 20 mM His pH 5.5 incubated at 25°C for seven days. A. Triplicates (red, green, blue) of the refolding curves of 2xSH Fc-C239i. B. Triplicates (red, green, blue) of the refolding curves of 2xCys Fc-C239i. C. Duplicates (red, green) of iDSB Fc-C239i.

WILEY-VCH

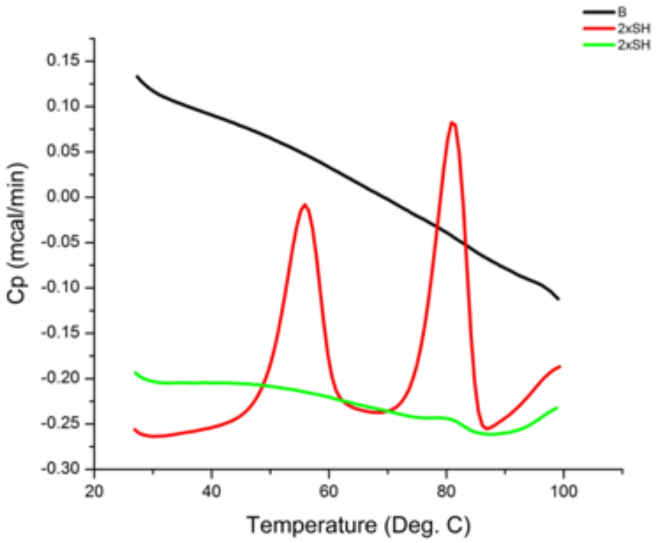


**Figure S. 8:** Raw data of the thermogram of the 2xSH Fc-C239i variant. The sample was heated a first time (red), cooled and reheated (green). The black line is the buffer baseline, 20 mM His pH 5.5.

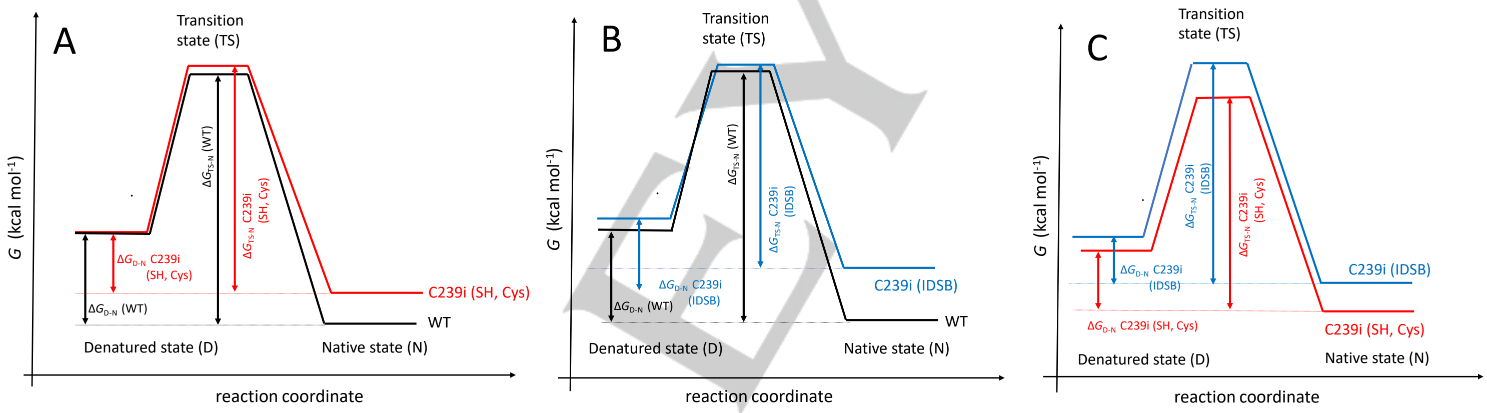


**Figure S. 9: Free energy diagrams combining the results from thermodynamic and kinetics stability experiments for the $C_H2$ domain of Fc-C239i variants and wild type. A. Free energy diagrams of 2xSH and 2xCys Fc-C239i vs wild type**. In this case, the energy of the native state (and to a lesser extent the TS) has increased as favourable interactions have been lost. The energy of the denatured state is the same. $\Delta G_{D\text{-}N}$ (WT) > $\Delta G_{D\text{-}N}$ C239i (SH, Cys), i.e. the WT is thermodynamically more stable than the inserted cysteine mutant in either the free thiol or cysteinylated forms. $\Delta G_{TS\text{-}N}$ (WT) > $\Delta G_{TS\text{-}N}$ C239i (SH, Cys), i.e., the rate constant for unfolding is smaller for WT than the inserted cysteine mutant in either the free thiol or cysteinylated forms and therefore the WT is more kinetically stable than the mutant. **B. Free energy diagrams of iDSB Fc-C239i vs wild type**. In this case, the energy of the native state (and to a lesser extent the TS) has increased as favourable interactions have been lost. The energy of the denatured state is higher as the additional disulfide bond restricts movement and decreases entropy. $\Delta G_{D\text{-}N}$ (WT) > $\Delta G_{D\text{-}N}$ C239i (IDSB), i.e. the WT is thermodynamically more stable than the inserted cysteine mutant in the IDSB form as formation of the IDSB greatly disrupts the structure and interactions in the native state to a much larger degree that it increases the energy of the denatured state by restricting entropy. $\Delta G_{TS\text{-}N}$ (WT) > $\Delta G_{TS\text{-}N}$ i239C (IDSB) i.e., the rate constant for unfolding is smaller for WT than the IDSB variant and therefore the WT is more kinetically stable than the mutant. **C. Free energy diagrams of iDSB Fc-C239i vs 2xSH and 2xCys Fc-C239i.** $\Delta G_{D\text{-}N}$ C239i (SH, Cys) > $\Delta G_{D\text{-}N}$ C239i (IDSB): C239i (SH, Cys) is more stable than the IDSB. Even though the introduction of an additional disulfide bridge stabilises the protein through increasing the energy of the denatured state, it destabilises the native to a much greater extent and is therefore overall very destabilising. $\Delta G_{TS\text{-}N}$ C239i (IDSB) > $\Delta G_{TS\text{-}N}$ C239i (SH, Cys), i.e., the rate constant for unfolding is smaller for C239i IDSB than the C239i (SH, Cys) and therefore the IDSB is more kinetically stable than the free thiol, cysteinylated forms even though it is less thermodynamically stable.

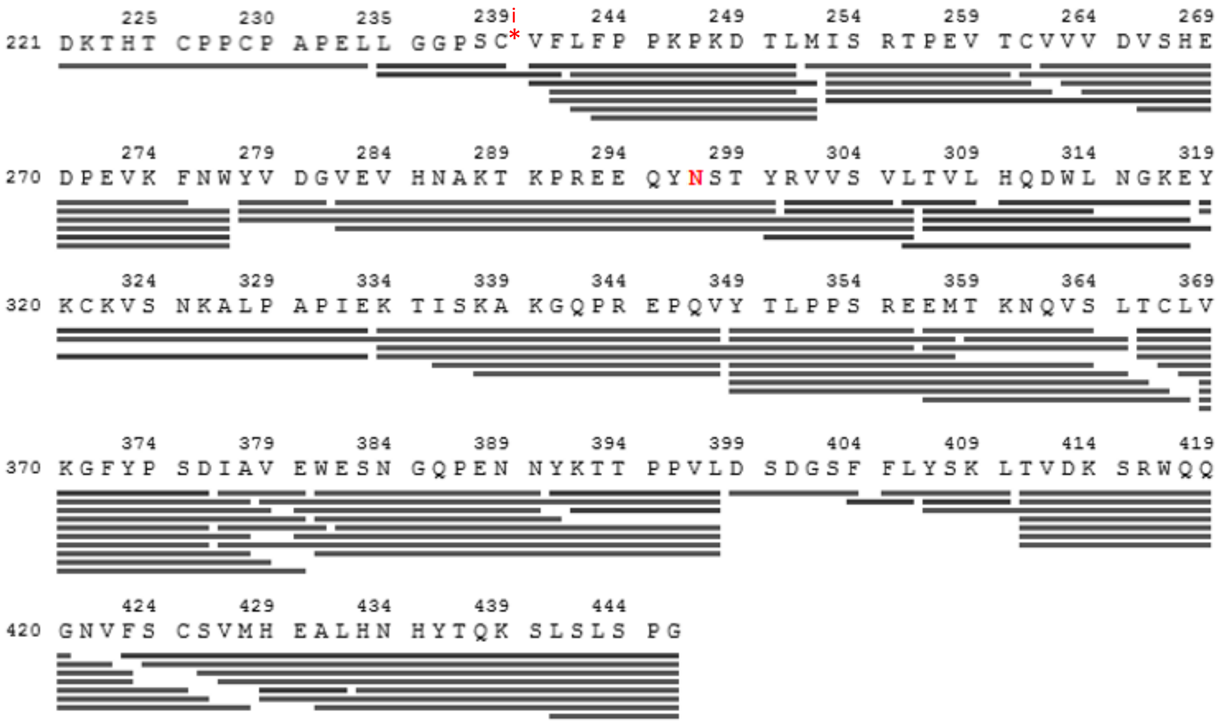


**Figure S. 10** : Peptide coverage of 2xCys Fc-C239i for millisecond HDX-MS experiments (100% coverage, 93 peptides, redundancy: 4.8). The red asterisk shows the location of the inserted cysteine. The canonical numbering of residues of an IgG1 was kept despite the insertion, therefore serine is at position 239 and the cysteine is 239i. Red N (Asn297) denotes N-glycan modification site.

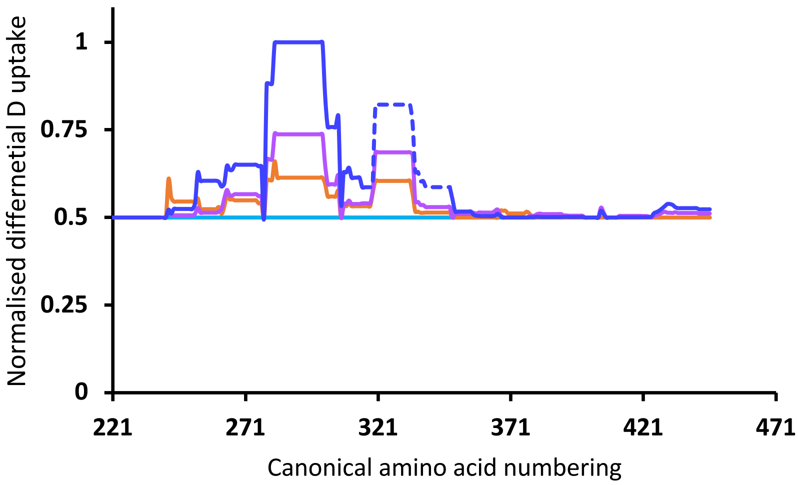


**Figure S. 11:** Normalised difference plot obtained after summing the significant D-incorporations relative to the wild-type. Light blue: wild-type (NIST mAb Fc) baseline. Orange: iDSB enriched Fc-C239i minus WT. Purple: 2xCys enriched Fc-C239i minus WT. Dark blue: iDSB enriched Fc-C239i minus WT. The dashed blue line corresponds to the region showing mixed EX1 and EX2 kinetics.

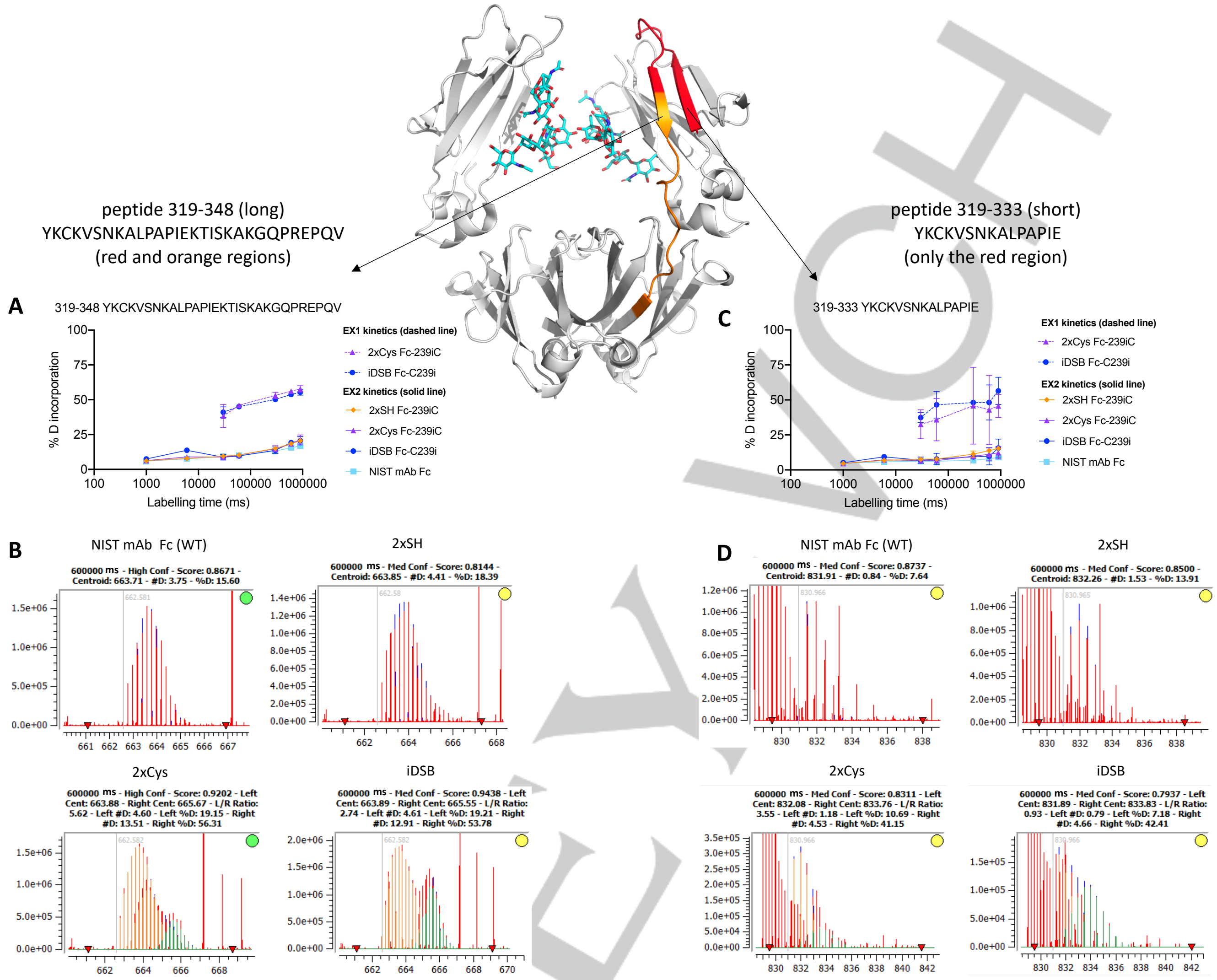


**Figure S. 12**: Details of mixed EX1/EX2 kinetics in the last two $\beta$-sheets of the $C_H2$ domain (colored on the crystal structure). **A.** Uptake plot representing the percentage in deuterium incorporation for the EX1 exchange (dashed line), and EX2 exchange (solid line) for the peptide 319-348 (YKCKVSNKALPAPIEKTISKAKGQPREPQV: red and yellow regions on the crystal structure). **B.** Raw spectra representing the unimodal or bimodal distributions for the deuterium uptake at 600 s. In the case of a bimodal distribution, the EX2 distribution is represented in orange and the EX1 distribution is in green. The EX1 distribution is three times more abundant for iDSB than for 2xCys at 600 s: the EX1 kinetics signal shifts by the same ratio as the iDSB residual amount in the 2xCys enriched sample (33.4%, **Table S. 1**), which supports the hypothesis that the the EX1 kinetics comes from the iDSB form. **C.** Uptake plot representing the percentage in deuterium incorporation for the EX1 exchange (dashed line), and EX2 exchange (solid line) for the peptide 319-333 (YKCKVSNKALPAPIE: red region on the crystal structure). **D.** Raw spectra representing the unimodal or bimodal distributions for the deuterium uptake at 600 s . More discussion in Supporting Information, section 4.2.

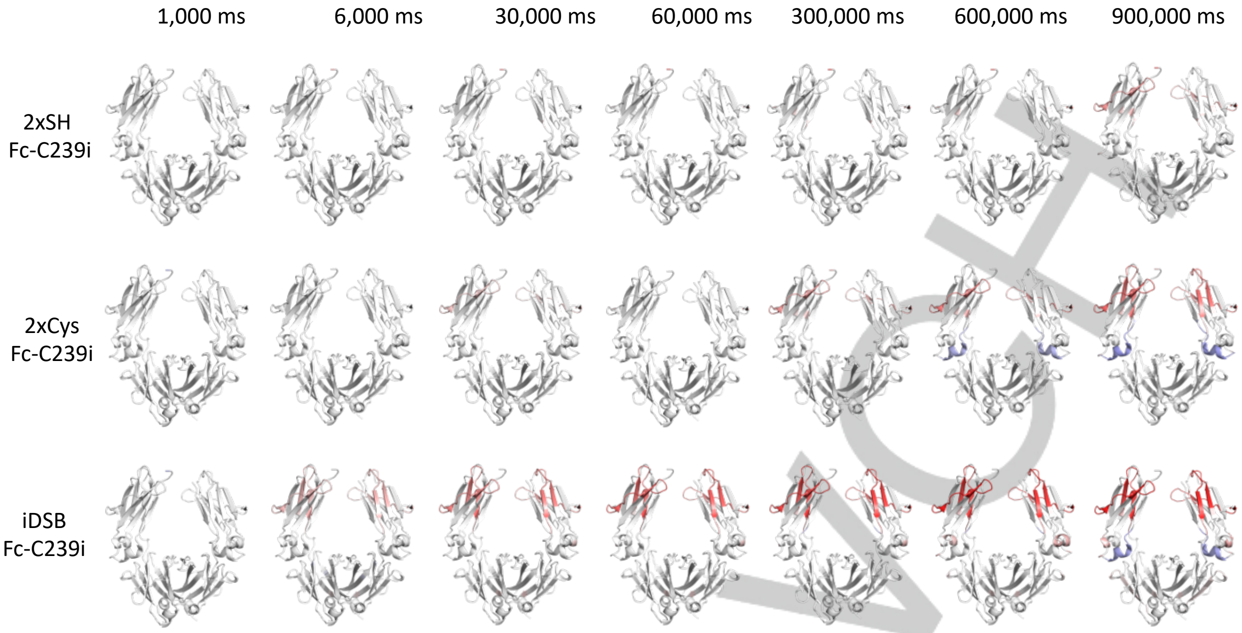


**Figure S. 13:** Crystal structure (3AVE) showing the changes in deuterium exchange, with the colour scale blue (less exchange), white, red (more exchange) per timepoint obtained with the second analysis method for the three enriched Fc-C239i variants compared to wild type. iDSB Fc-C239i is the variant that exchanges deuterium the most and each variant is normalized to this level.

### 3 Supplementary Tables

**Table S. 1:** Quality control of the Fc-239i variants after enrichment

| | 2xSH Fc-C239i (%) | 2xCys Fc-C239i (%) | iDSB Fc-C239i (%) |
|---|---|---|---|
| iDSB | 3.6 | 33.4 | 91.3 |
| 2xCys | 0.2 | 54.4 | 8.4 |
| 1xCys + 1xSH | 0.4 | 10.6 | 0.0 |
| 2xSH | 95.8 | 1.0 | 0.0 |
| 2x GSH | 0.0 | 0.3 | 0.1 |
| 1xGSH + 1xCys | 0.0 | 0.1 | 0.1 |
| 1xGSH + 1xSH | 0.0 | 0.0 | 0.0 |

**Table S. 2:** Effect of the incubation time at 25 °C and the concentration of chemical denaturant on the proportion of the inserted-cysteine states for each of the three initially enriched variants (2xSH Fc-C239i, 2xCys Fc-C239i, iDSB Fc-C239i)

| % | 2xSH Fc-C239i | | | | 2xCys Fc-C239i | | | | iDSB Fc-C239i | | | |
|---|---|---|---|---|---|---|---|---|---|---|---|---|
| Incubation time (days) | 0 days | 0 days | 7 days | 7 days | 0 days | 0 days | 7 days | 7 days | 0 days | 0 days | 7 days | 7 days |
| [GdmCl] (M) | 0 M | 3.5 M | 0 M | 3.5 M | 0 M | 3.5 M | 0 M | 3.5 M | 0 M | 3.5 M | 0 M | 3.5 M |
| iDSB | 4.7 | 19.8 | 8.3 | 99.7 | 39.8 | 55.1 | 43.9 | 81.1 | 97.0 | 97.5 | 97.4 | 99.7 |
| 2xCys | 0.0 | 0.0 | 0.0 | 0.0 | 46.8 | 42.6 | 45.3 | 18.9 | 2.6 | 2.5 | 2.6 | 0.3 |
| 2xSH | 95.3 | 80.2 | 91.7 | 0.3 | 1.0 | 0.6 | 0.8 | 0.0 | 0.0 | 0.0 | 0.0 | 0.0 |
| 1xCys | 0.0 | 0.0 | 0.0 | 0.0 | 12.3 | 1.7 | 10.0 | 0.0 | 0.4 | 0.0 | 0.0 | 0.0 |

**Table S. 3:** Effect of the concentration of denaturant after 7 days of incubation at 25 °C on 2xSH Fc-C239i enriched variant

| [GdmCl] (M) | iDSB (%) | | 2xCys (%) | | 1xCys + 1xSH (%) | | 2xSH (%) | | 2x GSH (%) | | 1xGSH + 1xCys (%) | | 1xGSH + 1xSH (%) | |
|---|---|---|---|---|---|---|---|---|---|---|---|---|---|---|
| | mean | SD | mean | SD | mean | SD | mean | SD | mean | SD | mean | SD | mean | SD |
| 0 | 2.7 | 0.01 | 0.080 | 0.003 | 0.41 | 0.03 | 96.80 | 0.02 | 0.0 | 0.0 | 0.0 | 0.0 | 0.0 | 0.0 |
| 0.5 | 24.5 | 0.4 | 0.6 | 0.01 | 0.272 | 0.002 | 74.5 | 0.4 | 0.0 | 0.0 | 0.07 | 0.02 | 0.0 | 0.0 |
| 1.0 | 86.6 | 0.4 | 1.8 | 0.1 | 0.0 | 0.0 | 11.3 | 0.5 | 0.0 | 0.0 | 0.25 | 0.04 | 0.0 | 0.0 |
| 1.5 | 92.05 | 0.02 | 1.5 | 0.1 | 0.0 | 0.0 | 6.3 | 0.1 | 0.0 | 0.0 | 0.0 | 0.0 | 0.0 | 0.0 |
| 2.0 | 96.3 | 0.1 | 1.8 | 0.1 | 0.0 | 0.0 | 1.8 | 0.1 | 0.0 | 0.0 | 0.06 | 0.03 | 0.0 | 0.0 |
| 2.5 | 96.9 | 0.2 | 2.0 | 0.1 | 0.0 | 0.0 | 0.9 | 0.1 | 0.1 | 0.0 | 0.0 | 0.0 | 0.0 | 0.0 |
| 3.0 | 98.1 | 0.1 | 1.3 | 0.1 | 0.0 | 0.0 | 0.56 | 0.01 | 0.0 | 0.0 | 0.0 | 0.0 | 0.0 | 0.0 |
| 3.5 | 97.3 | 0.7 | 2.4 | 0.7 | 0.0 | 0.0 | 0.32 | 0.01 | 0.0 | 0.0 | 0.0 | 0.0 | 0.0 | 0.0 |

**Table S. 4:** Effect of the concentration of denaturant after 7 days of incubation at 25 °C on 2xCys Fc-C239i enriched variant

| [GdmCl] (M) | iDSB (%) | | 2xCys (%) | | 1xCys + 1xSH (%) | | 2xSH (%) | | 2x GSH (%) | | 1xGSH + 1xCys (%) | | 1xGSH + 1xSH (%) | |
|---|---|---|---|---|---|---|---|---|---|---|---|---|---|---|
| | mean | SD | mean | SD | mean | SD | mean | SD | mean | SD | mean | SD | mean | SD |
| 0 | 31.3 | 0.4 | 54.9 | 0.2 | 12.6 | 0.2 | 1.11 | 0.00 | 0.08 | 0.01 | 0.0 | 0.0 | 0.0 | 0.0 |
| 0.5 | 42.3 | 0.2 | 52.9 | 0.1 | 4.2 | 0.2 | 0.46 | 0.02 | 0.0 | 0.0 | 0.0 | 0.0 | 0.0 | 0.0 |
| 1.0 | 64.2 | 0.8 | 35.8 | 0.9 | 0.0 | 0.0 | 0.0 | 0.0 | 0.0 | 0.0 | 0.0 | 0.0 | 0.0 | 0.0 |
| 1.5 | 69.5 | 0.1 | 30.5 | 0.1 | 0.0 | 0.0 | 0.0 | 0.0 | 0.0 | 0.0 | 0.0 | 0.0 | 0.0 | 0.0 |
| 2.0 | 78.2 | 0.6 | 21.8 | 0.6 | 0.0 | 0.0 | 0.0 | 0.0 | 0.0 | 0.0 | 0.0 | 0.0 | 0.0 | 0.0 |
| 2.5 | 82.7 | 0.3 | 17.2 | 0.3 | 0.0 | 0.0 | 0.0 | 0.0 | 0.0 | 0.0 | 0.0 | 0.0 | 0.0 | 0.0 |
| 3.0 | 77.71 | 0.02 | 22.26 | 0.01 | 0.0 | 0.0 | 0.0 | 0.0 | 0.0 | 0.0 | 0.0 | 0.0 | 0.0 | 0.0 |
| 3.5 | 78.6 | 0.6 | 21.4 | 0.6 | 0.0 | 0.0 | 0.0 | 0.0 | 0.0 | 0.0 | 0.0 | 0.0 | 0.0 | 0.0 |

**Table S. 5:** Effect of the incubation time in 3.5 M GdmCl at 25 °C on 2xSH Fc-C239i

| Incubation time (days) | iDSB (%) | | 2xCys (%) | | 1xCys + 1xSH (%) | | 2xSH (%) | | 2x GSH (%) | | 1xGSH + 1xCys (%) | | 1xGSH + 1xSH (%) | |
|---|---|---|---|---|---|---|---|---|---|---|---|---|---|---|
| | mean | SD | mean | SD | mean | SD | mean | SD | mean | SD | mean | SD | mean | SD |
| 0 | 10.6 | 0.3 | 0.3 | 0.0 | 0.1 | 0.0 | 89.0 | 0.3 | 0.0 | 0.0 | 0.1 | 0.0 | 0.0 | 0.0 |
| 1 | 96.5 | 0.2 | 1.9 | 0.1 | 0.0 | 0.0 | 1.6 | 0.1 | 0.0 | 0.0 | 0.0 | 0.0 | 0.0 | 0.0 |
| 2 | 97.5 | 0.1 | 1.5 | 0.0 | 0.0 | 0.0 | 1.0 | 0.0 | 0.0 | 0.0 | 0.0 | 0.0 | 0.0 | 0.0 |
| 3 | 97.3 | 0.6 | 1.8 | 0.6 | 0.0 | 0.0 | 0.8 | 0.0 | 0.0 | 0.0 | 0.0 | 0.0 | 0.0 | 0.0 |
| 4 | 97.3 | 0.2 | 1.9 | 0.1 | 0.0 | 0.0 | 0.8 | 0.1 | 0.0 | 0.0 | 0.0 | 0.0 | 0.0 | 0.0 |

**Table S. 6:** Effect of the incubation time in 3.5 M GdmCl at 25 °C on 2xCys Fc-C239i

| Incubation time (days) | iDSB (%) | | 2xCys (%) | | 1xCys + 1xSH (%) | | 2xSH (%) | | 2x GSH (%) | | 1xGSH + 1xCys (%) | | 1xGSH + 1xSH (%) | |
|---|---|---|---|---|---|---|---|---|---|---|---|---|---|---|
| | mean | SD | mean | SD | mean | SD | mean | SD | mean | SD | mean | SD | mean | SD |
| 0 | 41.4 | 0.3 | 55.6 | 0.4 | 1.80 | 0.02 | 1.0% | 0.0% | 0.09 | 0.01 | 0.0 | 0.0 | 0.0 | 0.0 |
| 2 | 58.1 | 0.1 | 41.7 | 0.1 | 0.0 | 0.0 | 0.0 | 0.0 | 0.09 | 0.01 | 0.0 | 0.0 | 0.0 | 0.0 |
| 3 | 66.2 | 0.1 | 33.7 | 0.1 | 0.0 | 0.0 | 0.0 | 0.0 | 0.0 | 0.0 | 0.0 | 0.0 | 0.0 | 0.0 |
| 4 | 69.6 | 0.1 | 30.36 | 0.03 | 0.0 | 0.0 | 0.0 | 0.0 | 0.1 | 0.1 | 0.0 | 0.0 | 0.0 | 0.0 |

WILEY-VCH

**Table S. 7:** Means of the melting temperatures of the antibody variants, run in triplicates.

| Antibody variant | $T_m1$ (°C) | $T_m2$ (°C) | $\Delta H1_{cal}$ (x $10^5$ kcal mol$^{-1}$) | $\Delta H2_{cal}$ (x $10^5$ kcal mol$^{-1}$) |
|---|---|---|---|---|
| NIST mAb Fc | 63.6 ± 0.2 | 79.7 ± 0.1 | 1.7 ± 0.5 | 1.4 ± 0.2 |
| 2xSH Fc-C239i | 55.4 ± 0.1 | 80.0 ± 0.3 | 1.3 ± 0.3 | 1.36 ± 0.06 |
| [1] 2xSH NEM capped Fc-C239i | 57.9 ± 0.2 | 80.1 ± 0.1 | 1.4 ± 0.3 | 1.27 ± 0.01 |
| 2xCys Fc-C239i | 56.6 ± 0.3 | 80.2 ± 0.1 | 1.3 ± 0.4 | 1.2 ± 0.1 |
| iDSB Fc-C239i | 54.8 ± 0.3 | 80.6 ± 0.2 | 0.5 ± 0.2 | 1.4 ± 0.1 |

[1]The Fc-C239i 2xSH NEM capped format was run just twice.
The errors are the standard deviations from repeated measurements.

**Table S. 8:** Unfolding kinetic parameters

| | $k_U^{H_2O}$ (s$^{-1}$) | $m_{k_U}$ |
|---|---|---|
| ln(k1) NIST mAb Fc | 0.5 ± 0.1 | 0.40 ± 0.04 |
| ln(k1) 2xSH | 3.6 ± 0.3 | 0.19 ± 0.01 |
| ln(k1) 2xCys | 2.6 ± 0.5 | 0.23 ± 0.03 |
| ln(k1) iDSB | 0.9 ± 0.1 | 0.34 ± 0.02 |
| ln(k2) NIST mAb Fc | (9 ± 4) E-06 | 1.69 ± 0.07 |
| ln(k2) 2xSH | (11 ± 4) E-06 | 1.67 ± 0.05 |
| ln(k2) 2xCys | (6 ± 4) E-06 | 1.8 ± 0.1 |
| ln(k2) iDSB | (8 ± 4) E-06 | 1.71 ± 0.08 |

The data points from **Error! Reference source not found.** C were fitted to the Equation 5 : $ln\, k_U^{[den]} = ln\, k_U^{H_2O} + m_{k_U}[den]$.

**Table S. 9**: Thermodynamic parameters fitted from the refolding curves

| | | $m_{I-N}$ (kcal mol$^{-1}$ M$^{-1}$) | | $[den]_{50\%\, I-N}$ (M) | | $\Delta G_{I-N}$ (kcal mol$^{-1}$) | | $m_{D-I}$ (kcal mol$^{-1}$ M$^{-1}$) | | $[den]_{50\%\, D-I}$ (M) | | $\Delta G_{D-I}$ (kcal mol$^{-1}$) | |
|---|---|---|---|---|---|---|---|---|---|---|---|---|---|
| 2xSH Fc-C239i | unfolding | 2.5 | 0.2 | 0.58 | 0.02 | 1.5 | 0.2 | 3.1 | 0.1 | 2.05 | 0.04 | 6.4 | 0.3 |
| | refolding | 1.4 | 0.3 | 0.52 | 0.03 | 0.7 | 0.2 | 3.0 | 0.1 | 1.98 | 0.03 | 5.9 | 0.3 |
| 2xCys Fc-C239i | unfolding | 2.099 | 0.003 | 0.9060 | 0.0001 | 1.902 | 0.002 | 5.3 | 0.6 | 2.30 | 0.01 | 12 | 1 |
| | refolding | 1.94 | 0.01 | 0.96 | 0.08 | 1.9 | 0.2 | 6.3 | 0.1 | 2.28 | 0.03 | 14.4 | 0.3 |
| iDSB Fc-C239i | unfolding | 1.3 | 0.1 | 0.75 | 0.05 | 1.0 | 0.1 | 4.5 | 0.2 | 2.272 | 0.002 | 10.3 | 0.5 |
| | refolding | 1.0 | 0.2 | 0.6 | 0.08 | 0.7 | 0.1 | 5.5 | 0.3 | 2.31 | 0.02 | 12.8 | 0.8 |
| NIST mAb Fc | unfolding | 3.2 | 0.1 | 1.69 | 0.02 | 5.5 | 0.2 | 3.9 | 0.5 | 2.25 | 0.04 | 9 | 1 |
| | refolding | 3.0 | 0.1 | 1.69 | 0.01 | 5.0 | 0.2 | 3.2 | 0.4 | 2.24 | 0.02 | 7.1 | 0.8 |

The values are single runs (NIST mAb Fc refolding), means of duplicates (2xCys Fc-C239i, iDSB Fc-C239i unfolding and refolding) and triplicates (2xSH Fc-C239i unfolding and refolding, 2xCys Fc-C239i refolding, NIST mAb Fc unfolding) and the standard deviation of the repeats.

**Table S. 10:** Summary of HDX mass spectrometry experimental details

| | |
|---|---|
| Dataset | 2xSH Fc-C239i (5 µM), 2xCys Fc-C239i (5 µM), iDSB Fc-C239i (5 µM), NIST mAb Fc (5 µM) |
| HDX reaction details | Equilibration in $H_2O$: 20 mM His buffer pH 5.5<br>Labelling in $D_2O$: 20 mM His buffer pD 5.5<br>Both were done at 20 °C |
| HDX time course | 1000, 6000, 30000, 60000, 300000, 600000 and 900000 ms |
| HDX controls | 0 s |
| Number of peptides | 93 |
| Sequence coverage | 100 % |
| Redundancy | 4.8 |
| Average peptide length | 12.5 AA |
| Replicates | Technical replicates: 3 |
| Repeatability | Average Student's t-distribution 95% confidence interval for all peptide and all time points for the WT (NIST mAb Fc):<br>#D: 0.080403<br>%D: 0.90323 |
| Significant differences in HDX | The incorporation of deuterium per peptide is significant compared to the wild-type if p-value < 0.05 (t-test) |

### 4 Supplementary Results

#### 4.1 Verification of the interchain disulfide bridge

Fc-C239i enriched in the iDSB form was digested with the Lys C enzyme and the digested peptides were further fragmented inside the mass spectrometer by collision induced dissociation (CID). The aim of this experiment is to focus on the peptide corresponding to the hinge (**Figure S. 4 A**), which contains the two canonical cysteines as well as the inserted one at the position 239 all forming disulfide bridges, and to fragment it by CID to find evidence showing that the additional interchain disulfide bridge is indeed linking the two heavy chains, parallel to the other two. The peptide in the hinge, corresponding to a +4 ion (1415.17 m/z) was isolated and fragmented, and the obtained peptides were identified. Peptides linked by a disulfide bridge between the inserted cysteines 239 were identified and the amino acid sequence was reconstituted amino acid by amino acid (**Figure S. 4 B**): they are composed of a neutral peptide truncated at the N-terminus only (resulting from a fragmentation that generated a y ion and a neutral peptide, as the charges were retained on the N-terminus fragmented peptide[35]), and a + 1 charged peptide truncated both at the N-terminus, as a neutral fragment (same explanation as above), as well as at the C-terminus as a b ion (after the fragmentation event, the charge was retained at the N-terminus). These findings demonstrate that the additional disulfide bridge is indeed an interchain disulfide bridge between the two 239 inserted cysteine linking the two heavy chains, as the canonical disulfide bridges do.

#### 4.2 Chemical denaturation curves

##### 4.2.1 Reproducibility

The chemical denaturation curves of the control NIST mAb Fc, Fc-C239i free-thiol and cysteinylated were performed in triplicates, and those of the Fc-C239i iDSB were performed in duplicates. The thermodynamic parameters obtained for each are very similar or within error (**Table S. 9**), which shows that the data are highly reproducible (**Figure S. 6 A-C**).

##### 4.2.2 Reversibility

The reversibility of chemical denaturation of the Fc constructs was investigated by undertaking measurement of refolding curves, where the protein has been unfolded at the start, then diluted into a range of denaturant concentrations and finally measured once equilibrium has been obtained.
For Fc-C239i enriched in the free thiol form, the $m$-value of the first unfolding transition between the native and intermediate states ($m_{I-N}$) is quite different to that obtained from the unfolding curve data. This is due to the fact that there is no native baseline in the refolding curves and data collection starts at 0.4 M GdmCl, meaning it is difficult to get an accurate m value. Nevertheless, the denaturation midpoints are very similar, which establishes unfolding is reversible. For the other mutants, the thermodynamic parameters are within error (**Table S. 9**), which proves that the denaturation is reversible and at equilibrium for these time points. Replicate refolding denaturation curves be found in **Figure S. 7**.

#### 4.3 Discussion on accuracy of measurements from chemical denaturation curves and differential scanning calorimetry

It is important to keep in mind that the enriched states 2xSH and 2xCys convert to iDSB if there is no NEM, and therefore the chemical denaturation curves, as well as the thermal stability experiments most probably, have a gradient of variants as the concentration of denaturant or temperature increases, starting from almost totally 2xSH or 2xCys, and tending towards iDSB during the denaturation. The only data sets with exactly the same material all along the experiments are those with the iDSB enriched variant and the 2xSH NEM capped.

#### 4.4 Discussion on mixed EX1/EX2 kinetics in lower $C_H2$ domain

The peptide 319-348 going from the β-strand F until the beginning of the $C_H3$ domain shows a very clear and strong EX1 kinetic signal for iDSB Fc-C239i enriched variant, whereas the 2xCys Fc-C239i enriched variant just shows a smaller signal for the EX1 kinetics, equivalent to 1/3 of the iDSB's intensity (green distribution, **Figure S. 11 B**). The EX1 kinetics signal shifts by the same ratio as the iDSB residual amount in the 2xCys enriched sample (33.4%, **Table S. 1**), which supports the hypothesis that the the EX1 kinetics comes from the iDSB form. EX1 kinetics were also observed in the shorter overlapping peptide 319-333 (**Figure S. 12 C**), but not in the other overlapping peptide 334-348 nor in adjacent regions. This suggests that the EX1 deuterium exchange kinetics come from the antiparallel β-strands F and G (**Figure 3 C**) which are undergoing local unfolding, i.e. unzipping and rezipping without breaking the surrounding H-bonds of the β-sheet. The mixed EX1/EX2 kinetics suggest that there is a mixed population of folded and unfolded forms of the $C_H2$ domain for the iDSB variant ($iDSB_D \rightleftharpoons iDSB_N$).

#### 4.5 Comparison between two HDX-MS processing methods

The results from both methods (section 1.9, **Figure 3** and **Figure S. 13**) agree on the regions of greatest deuterium exchange, which occur in the β-sheet composed of the strands C, F and G (green β -sheet, **Figure 3 C**) as well as with the β-strand D. There are two regions where one analysis method do not show significant change, and while statistically significant represents a small difference in fractional exchange. The first difference is located in the upper $C_H2$ domain: the first analysis method shows significant exchange at the beginning of the $C_H2$ domain, specifically between the residues Ser 254 to Trp 277 involved in two β-strands B and C (**Figure 3 C**), but of low amplitude; the second method does not show any significant change in that region. The second difference involved a loop between strands A and B, which show significant low protection according to the second processing method, but no significant protection with the first method. Overall, both analysis methods are in broad agreement with the differences highlighting that different approaches in calculating fractional exchange across timepoints and the statistical method employed to determine significance will result in slightly different interpretation of a common data set.

WILEY-VCH
